# Spatially informed phenotyping by cyclic-in-situ-hybridization identifies novel fibroblast populations and their pathogenic niches in systemic sclerosis

**DOI:** 10.1101/2024.12.28.630505

**Authors:** Yi-Nan Li, Tim Filla, Andrea-Hermina Györfi, Minrui Liang, Veda Devakumar, Alexandru Micu, Hongtao Chai, Christina Bergmann, Ann-Christin Pecher, Jörg Henes, Pia Moinzadeh, Thomas Krieg, Alexander Kreuter, Georg Schett, Bernhard Homey, Sascha Dietrich, Jörg H.W. Distler, Alexandru-Emil Matei

## Abstract

Spatially non-resolved transcriptomic data identified functionally distinct populations of fibroblasts in health and disease. However, in-depth transcriptional profiling *in situ* at single-cell resolution has not been possible so far. Here, we studied fibroblast populations in the skin of SSc patients and healthy individuals using cyclic in situ hybridization (cISH) as a novel approach for spatially-resolved transcriptional phenotyping with subcellular resolution. cISH deconvoluted the heterogeneity of 20,979 cells including 3,764 fibroblasts (FB). BANKSY-based spatially-informed clustering identified nine FB subpopulations, with SFRP2+ RetD FB and CCL19+ nonPV FB as novel subpopulations that reside in specific cellular niches and display unique gene expression profiles. SFRP2+ RetD FB and CCL19+ nonPV FB as well as COL8A1+ FB, display altered frequencies in SSc skin and play specific, disease-promoting roles for extracellular matrix release and leukocyte recruitment as revealed by their transcriptional profile, their cellular interactions and ligand-receptor analyses. The frequencies of COL8A1+ FB and their interactions with monocytic cells and B cells are associated with progression of skin fibrosis in SSc. In summary, our spatially-resolved transcriptomic approach identified novel fibroblast subpopulations deregulated in SSc skin with specific pathogenic roles, some of which may potentially serve as biomarkers for progression of skin fibrosis.

## Introduction

Systemic sclerosis (SSc) is a connective disease characterized by autoimmunity and inflammation, which promote extensive fibrotic remodeling of the skin and various internal organs (1). The progressive accumulation of extracellular matrix in affected tissues impairs organ function and commonly leads to organ failure, resulting in high morbidity and mortality of SSc patients (2). Fibroblasts are the key effector cells for release of extracellular matrix in fibrosis (2–5). However, fibroblasts are a heterogenous population of cells with specific, functionally distinct subpopulations. Understanding fibroblast heterogeneity, particularly in the context of specific diseases, is still in its early stages. Moreover, we are lacking information on the organization of fibroblast subpopulations *in vivo* and the intricate cellular interactions that occur within their native tissue environment.

Single-cell omics allows for the study of cellular subpopulations at previously unattainable resolution. So far, non-spatially resolved single-cell RNA sequencing (scRNA-seq) has been predominantly used to phenotype cellular changes in rheumatic diseases. scRNA-seq revealed that fibroblasts constitute a heterogenous population of cells with distinct transcriptional patterns and functions (6). scRNA-seq data from SSc skin revealed between six to ten distinct subpopulations of fibroblasts (7–9). However, scRNA-seq, like other existing single cell techniques, requires tissue disaggregation and single cell separation. As a result, these approaches cannot provide information on the spatial localization of cells inside the tissue, nor can they characterize tissue niches and the cellular interplay that occurs within them. Furthermore, recovery rates can vary considerably amongst cell populations, thus influencing quantification results. Spatial sequencing approaches overcome the lack of spatial context, but approaches published so far have a modest resolution of around 100 µm and are thus not able to resolve individual cells (7).

Here, we aimed to overcome these limitations using cyclic in-situ hybridization as a spatial transcriptomic approach with subcellular resolution of 120 nm. Using a cISH panel customized for studies of fibroblasts by enrichment of fibroblast marker genes, we classify and quantify fibroblast subpopulations based on their transcriptional and spatial context, characterize their transcriptional features, their microenvironment and their cellular interactions and the predictive power of these molecular features for progression of skin fibrosis in SSc.

## Results

### Study design and identification of major cell types in SSc and control skin by cISH

To characterize fibroblast subpopulations in SSc skin, their microenvironment and their ability to predict progression of fibrosis in SSc, we performed cISH from skin samples of SSc patients enriched for early, diffuse cutaneous involvement, with clinically active disease and sex-matched and age-matched healthy controls (Figure 1A). To increase the resolution of our gene panel for fibroblast subpopulations, we extended the standard panel of 950 genes with 86 additional probes targeting 19 marker genes of fibroblast subpopulations as described in previous single cell OMICs studies (Figure S1, Table S2) (6, 8). We recovered a total of 20,979 cells (patients with SSc: 9,876 cells; healthy individuals: 11,103 cells) (Figure 1A-C). These single cells included 3,764 fibroblasts, identified by their high expression levels of *COL1A1*, *COL1A2*, *COL3A1* and *DCN* (Figure 1B, D). We further identified other major cell populations from human skin, including keratinocytes (expressing *KRT1*, *KRT10*, *KRT14*, and *KRT15*), immune cells (expressing the pan-leukocyte gene *PTPRC* and specific genes for immune cells from the lymphoid and myeloid lineages such as *CD4*, *CD68*, *CD163* and *LYZ*), endothelial cells (expressing *PECAM1*, *CDH5*, *ENG* and *VWF*) and vascular smooth muscle cells (expressing *TAGLN*, *ACTA2* and *MYL9*) (Figure 1D). These cell types had the expected spatial distribution in human skin, further validating their identity (Figure 1E).

**Figure 1:**
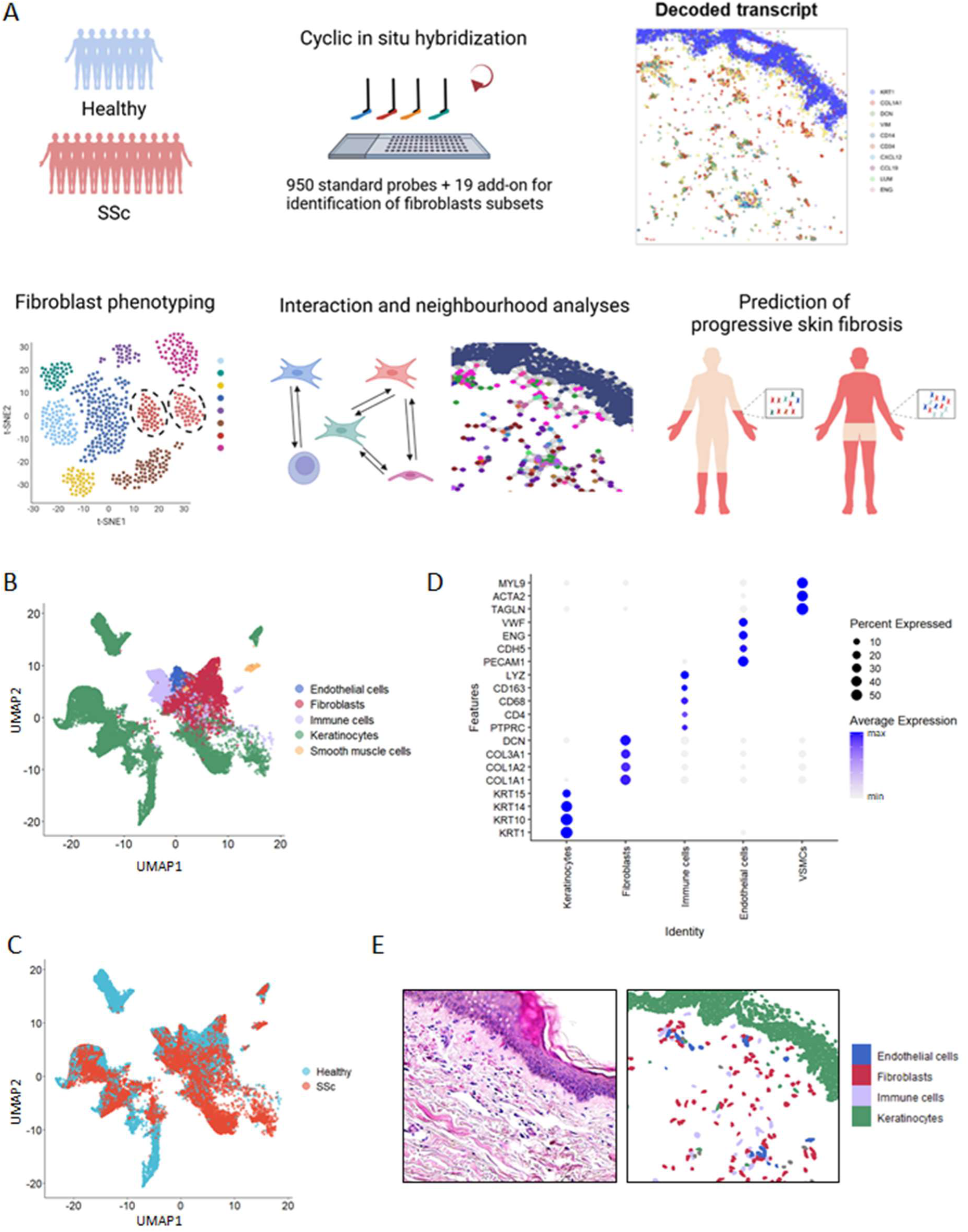
Characterization of skin tissue organization in SSc using cISH. (A) Schematic overview of the experimental and analytic workflow. The transcriptomic profiles of skin tissue from healthy and SSc donors were analyzed by imaging-based spatial transcriptomics using cISH. The cell populations in the skin were characterized based on the transcriptional profile. Taking advantage of spatial information, we performed cellular interaction analysis and evaluated whether changes in cell frequencies or spatial relationships might predict progression of skin fibrosis. (B) UMAP plot showing 20,979 cells detected in the skin samples. The plot is colored by the major cell types within the skin tissue. (C) UMAP plot showing the cells detected in the skin samples obtained from healthy donors and SSc patients. (D) Dot plot indicating marker gene expression of the major cell populations in the skin tissue. The size of the dot represents the percentage of the cells that expressed the specified gene. The dots were colored by levels of the expression of specified gene. (E) Representative images of HE staining and distribution of cISH-detected cells in the skin tissue. This figure was created with BioRender.com. cISH, cyclic in situ hybridization; SSc, systemic sclerosis; UMAP, Uniform Manifold Approximation and Projection; HE, hematoxylin and eosin.

### Spatially informed phenotyping of fibroblasts in SSc and control skin identifies new fibroblast subpopulations

Conventional subclustering of the fibroblasts based on their gene expression revealed seven subpopulations (Figure S2A). We annotated these fibroblast subpopulations based on their marker expression: SFRP2+ FB, COL8A1+ FB, PI16+ FB, CCL19+ FB, ACKR3+ FB, GSN+ FB, and DUSP1+ PB (Figure S2B, C). SFRP2+, COL8A1+, CCL19+ and PI16+ FB were described in previous scRNA-seq studies (6, 8–10), highlighting that cISH-based phenotyping with the extended panel could reliably identify key fibroblasts subsets in the skin.

We further aimed to refine the fibroblast phenotyping and performed spatially informed subclustering of fibroblasts using the BANKSY method. This algorithm leverages the spatial information of the dataset to identify the neighbors of each cell and perform clustering based not only on the own gene expression, but also on the average expression and expression gradients among each cell’s neighbors (Figure 2A) (11).

**Figure 2:**
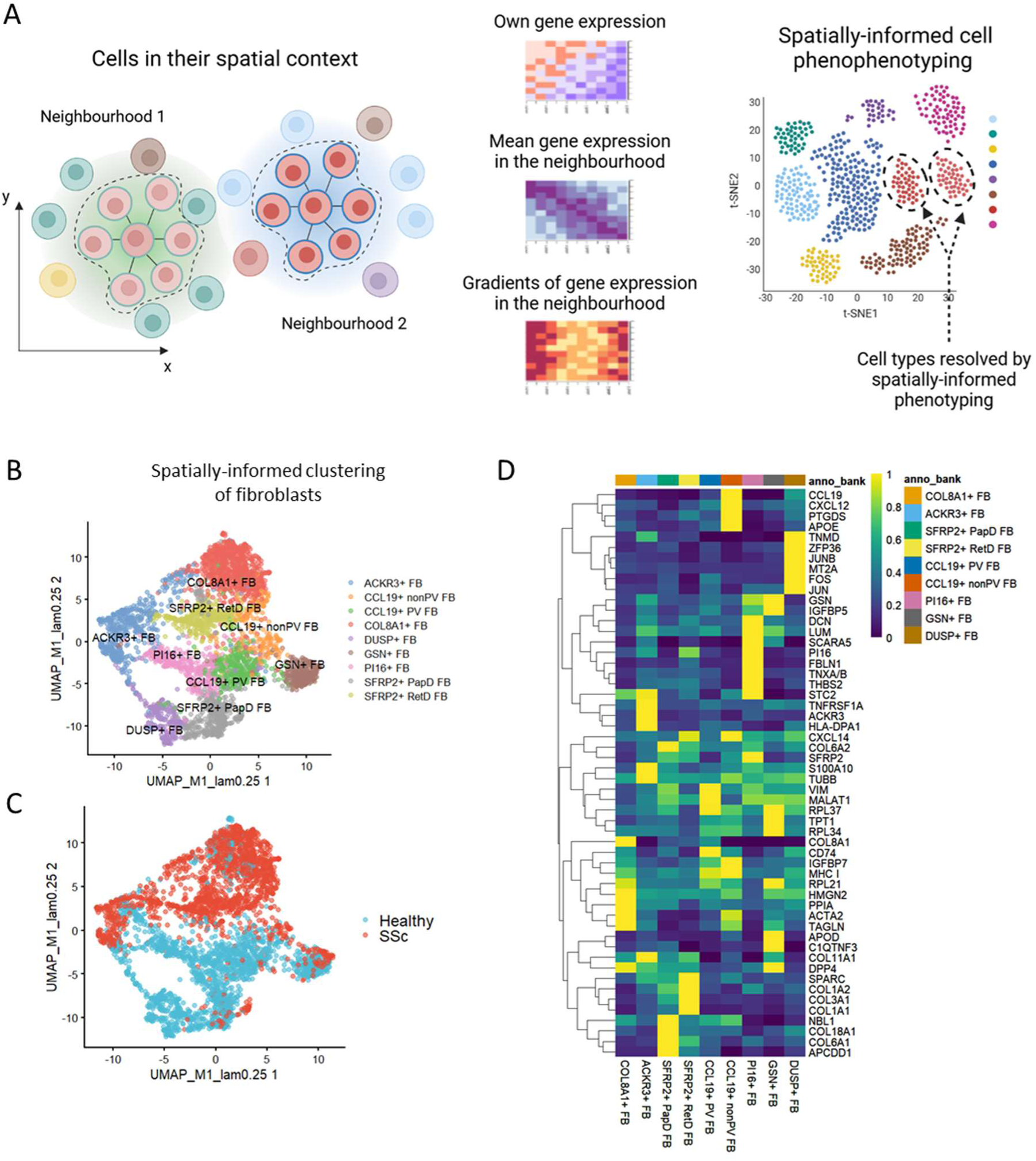
Fibroblasts populations identified by spatially informed clustering. (A) An illustration showing the workflow for spatially informed cell phenotyping by BANKSY-based spatial clustering. In addition to the expression matrix of the cell of interest, the transcriptional profiles of the local environment, including averaged expression and the expression gradient of neighboring cells, were computed by BANKSY and subjected to phenotyping by clustering. (B) UMAP showing fibroblast populations identified by spatially informed clustering using BANKSY. (C) UMAP showing the BANSY-identified populations from healthy and SSc donors. (D) Expression heatmap of the marker genes for the BANKSY-identified populations. This figure was created with BioRender.com. BANKSY, Building Aggregates with a Neighborhood Kernel and Spatial Yardstick; UMAP, Uniform Manifold Approximation and Projection; SSc, systemic sclerosis.

Visualization of the BANKSY-based clustering by dimensionality reduction with the algorithm uniform manifold approximation and projection (UMAP) demonstrated distinct patterns for fibroblasts derived from skin samples of patients with SSc and healthy individuals, indicative of major changes in the spatially informed phenotype of fibroblast populations in SSc (Figure 2B, C).

Using this approach, we identified nine subpopulations of fibroblasts and annotated them based on the correspondence of the BANKSY-based clusters with the conventional (non-spatial) clusters, on the differential marker expression and on the spatial localization (Figure 2D and Figure S3C). Two BANKSY-based clusters corresponded to the SFRP2+ FB (defined by non-spatial clustering) (Figure S3A-C). Indeed, one of the two SFRP2+ populations was located in spatial proximity to the epidermal layer and was thus termed SFRP2+ PapD (papillary dermis) FB. In contrast, the other SFRP2+ population was located deeper in the dermis and was thus termed SFRP2+ RetD (reticular dermis) FB (Figure S3D). The average expression and expression gradients of the keratinocyte markers *KRT14* and *KRT5* were higher among the neighboring cells of SFRP2+ FB RetD than those of the SFRP2+ FB PapD (Figure S3F), further confirming their distinct spatial distribution.

Similarly to the SFRP2+ FB, BANKSY-based clustering identified two subpopulations of the previously described CCL19+ FB cluster as defined by non-spatial clustering (Figure S3A-C). One of the CCL19+ FB, that we termed CCL19+ PV (perivascular) FB, was located around endothelial cells and consistently had a higher expression of the endothelial cell marker *PECAM1* and the pericyte marker *RGS5* among its neighbors than the CCL19+ nonPV (non-perivascular) FB (Figure S3E, G). The latter subpopulation was more diffusely distributed in the dermis without enrichment in a particular dermal compartment (Figure S3E).

Of particular note, the SFRP2+ RetD and the CCL19 PV FB are not only distinct from the SFRP2+ PapD and the CCL19 nonPV FB, respectively, with regards to their spatial location, but also with regards to their own gene expression, with e.g. higher levels of *COL1A1* and *COL3A1* expressed by the SFRP2+ RetD than SFRP2+ PapD FB, and higher levels of *CXCL12* and *PTGDS* expressed by the CCL19+ nonPV than the CCL19+ PV FB (Figure 2D).

In contrast, the other five subpopulations of fibroblasts defined by non-spatial clustering were homogenous not only regarding their own gene expression, but also to their spatial distribution and were not further subclustered by the BANKSY method (Figure S3A-C).

Integration with two scRNA-seq datasets, i.e. GSE138669 (6) and GSE195452 (8), demonstrated that neither of the label transfer-based annotations from the scRNA-seq to the cISH dataset could reliably distinguish between the SFRP2+ PapD and RetD FB, or between the CCL19+ PapD and RetD FB, respectively (Figure S4, Figure S5), highlighting that the spatially informed BANKSY-based annotation can refine the fibroblast phenotyping performed with previous methods.

To further validate the distinct spatial relationship of the SFRP2+ PapD and RetD FB with regards to epidermis, and of the CCL19+ PV and CCL19+ nonPV FB with regards to vessels, we analyzed a complementary spatial transcriptomic dataset. While this second dataset did not provide single-cell resolution, it provided a higher sequencing depth that covers the whole transcriptome. Since each spot contains approximately four cells (12), we deconvoluted the composition of spots by the Seurat anchor-based reference mapping (13). Even though this deconvolution method relies exclusively on the own gene expression of each cell, we could indeed show that spots enriched in SFRP2+ PapD FB, in contrast to those enriched in SFRP2+RetD FB, were located predominantly in the papillary dermal region (Figure S6A). Similarly, spots enriched in CCL19+ PV FB were located predominantly in the perivascular region, whereas those enriched in CCL19+ nonPV FB demonstrated a diffuse localization in the dermis (Figure S6B).

### Changes in frequencies of fibroblast subpopulations in SSc skin

In line with previous studies, the total number of fibroblasts was similar in SSc compared to control skin (1658 vs. 1732, respectively) (14). However, SFRP2+ RetD FB and the COL8A1+ FB had significantly higher frequencies in SSc, whereas the PI16+ FB, the DUSP+ FB and the CCL19+ PV FB had modestly lower frequencies in SSc (Figure 3). The changes in frequencies of these subsets in SSc are consistent with the increases of SFRP2+ FB and COL8A1+ FB, and decrease in PI16+ FB in SSc skin reported by scRNA-seq-based studies (6–9). Of note, the populations resolved only by BANKSY clustering, i.e. the SFRP2+ RetD and PapD FB, and CCL19+ PV and nonPV FB, respectively, demonstrated distinct changes in SSc. SFRP2+ PapD FB were numerically decreased in frequency in SSc, whereas the SFRP2+ RetD FB were increased, and CCL19+ nonPV were numerically increased in SSc, whereas the CCL19+ PV were decreased (Figure 3A, C, E).

**Figure 3:**
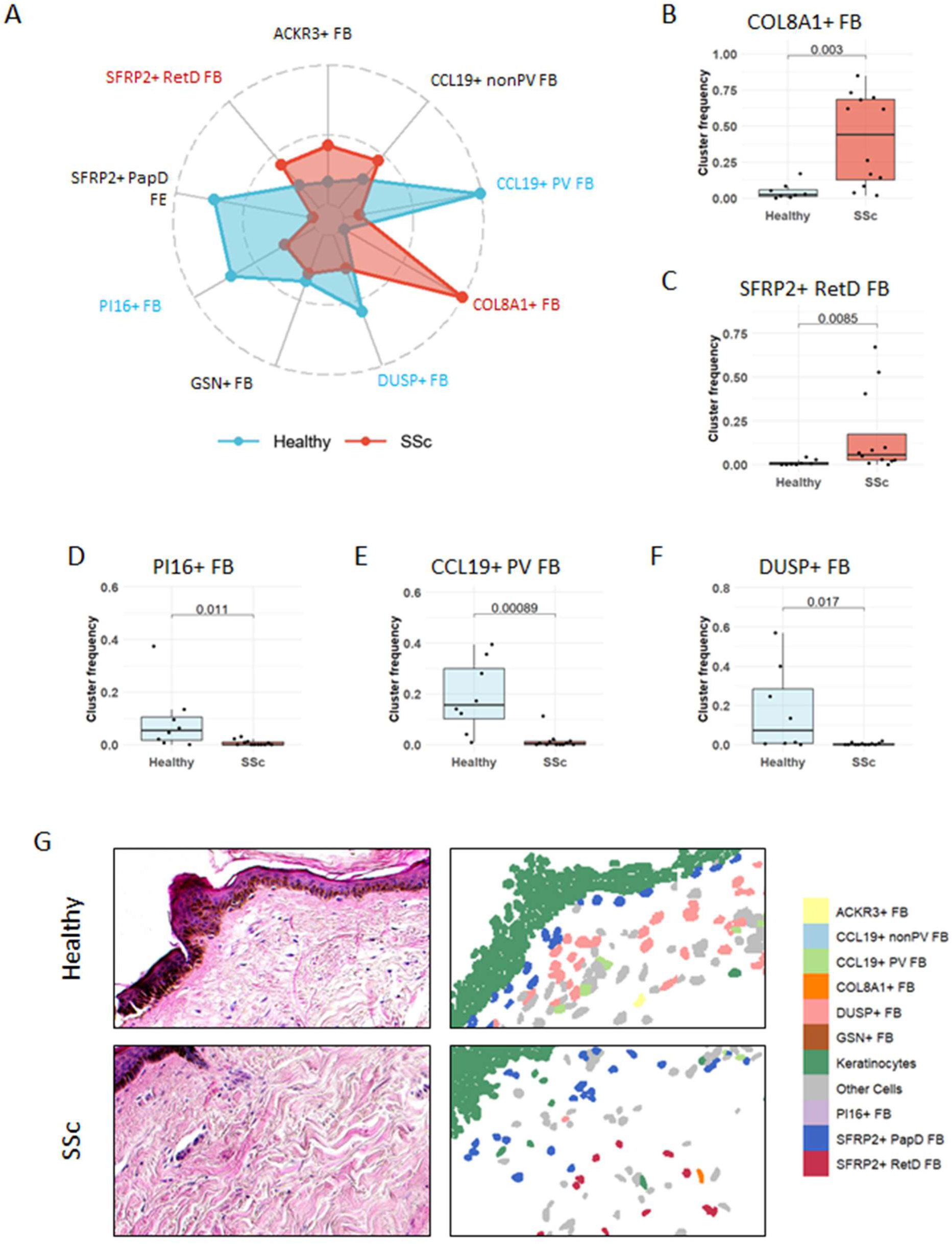
Changes in frequencies of fibroblast subsets identified by spatially informed phenotyping in SSc. (A) An overview of the changes in the abundance of BANKSY-identified fibroblast populations. The abundance was quantified by the proportion of the specified population within the fibroblast component. Z-normalized median proportions were plotted on the radar chart. The statistically significant changes were highlighted in red (increased in SSc) and blue (increased in healthy). (B-F) Changes in frequencies of COL8A1+ FB (B), SFRP2+ RetD FB (C), PI16+ FB (D), CCL19+ PV FB (E) and DUSP+ FB (F) in SSc compared to control skin. (G) Representative images of BANKSY-identified populations in the skin tissue, along with HE stainings of the corresponding regions. The statistical significance was determined by the Mann-Whitney U-test. P-values below 0.05 were considered significant. BANKSY, Building Aggregates with a Neighborhood Kernel and Spatial Yardstick; SSc, systemic sclerosis; HE, hematoxylin and eosin.

We next performed pseudotime analysis on the spatially informed fibroblast subsets using Monocle (15). We defined as root (starting point) of the pseudotime the PI16+ FB, since they were previously described as a fibroblast subset with stem cell properties that can differentiate into specialized fibroblasts (16, 17), and that in our, as well as in publicly available scRNA-seq dataset have a lower frequency in SSc (6). The pseudotime analysis revealed three differentiation trajectories, one going through the SFRP2+ PapD FB and ending on DUSP+ FB, one going through CCL19+ PV, then through CCL19+ nonPV FB and ending on the GSN+ FB, and a third one going through ACKR3+ FB, followed by SFRP2+ RetD FB and ending on COL8A1+ FB (Figure S7). The latter differentiation trajectory started from a fibroblast subset with decreased frequency in SSc and ended in two subsets that are highly enriched in SSc, which suggests that the fibroblast subsets enriched in SSc are terminally differentiated from precursor fibroblasts (among other cell types), in line with previous publications (16–18). Of note, while the CCL19+ PV FB and CCL19+ nonPV FB subsets aligned along one trajectory, and are thus likely developmentally related, the SFRP2+ PapD FB and the SFRP2+ RetD FB were located along two distinct trajectories. This suggests that fibroblast phenotyping by marker expression alone might be insufficient for evaluation of differentiation trajectories between the fibroblast subsets, and that fibroblasts with partially overlapping marker expression profile, but with distinct spatial locations, might have distinct developmental cues.

### Functional heterogeneity of the spatially informed fibroblast subsets

We next aimed to evaluate whether the fibroblast subsets identified by spatially informed phenotyping are functionally distinct. We first computed an ECM module score, as well as scores for different ECM elements, i.e. collagens, glycoproteins and proteoglycans, with the gene sets curated in the Matrisome project (Figure 4A and Figure S8A) (19). All fibroblast subsets had higher ECM and collagens scores than the other cell types, re-confirming their identity (Figure 4A). The SFRP2+ RetD had the highest ECM score and collagen score of all fibroblast subpopulations (Figures 4A), while the PI16+ FB had the highest glycoproteins and proteoglycans scores (Figure S8A), pointing to specific roles in ECM production for the different subsets. The TGFβ pathway activity inferred by using PROGENy signatures (20) was most enriched in SFRP2+ RetD FB (Figure 4B). Comparison of fibroblast populations in SSc and normal skin revealed higher ECM and collagens scores and of TGFβ activity for several fibroblast populations in SSc skin (Figure S8B). Thus, not only changes in the frequencies of fibroblast populations but also changes in the activation status of individual populations further shift the balance in SSc skin towards fibrotic remodeling.

**Figure 4:**
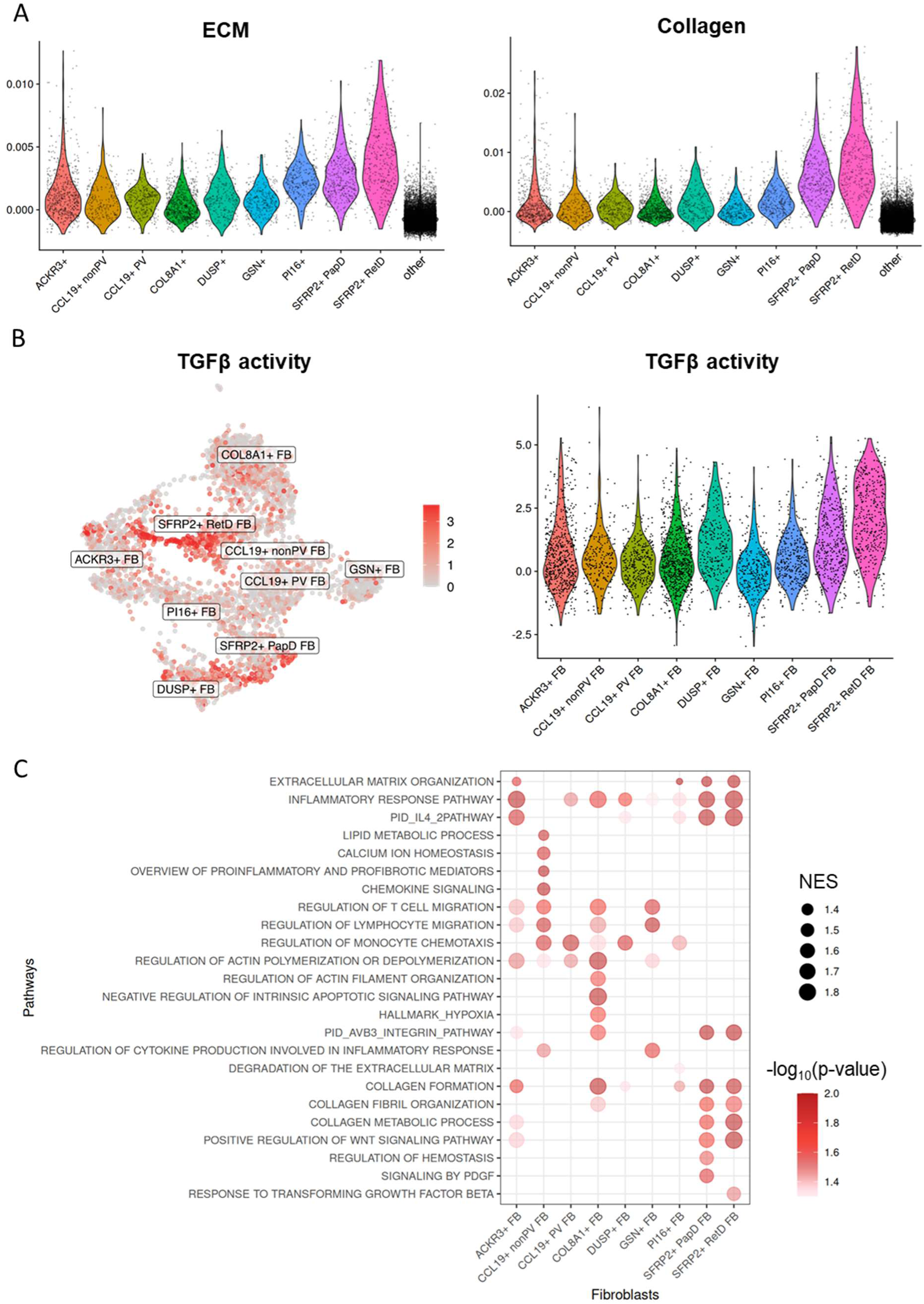
Distinct biological functions of spatially identified fibroblast populations. (A) Violin plots showing the ECM module scores of components obtained from the Matrisome project across fibroblast populations. Non-fibroblast populations were labeled as ‘other’ in the violin plots. (B) UMAP and violin plots illustrating TGFβ activities inferred using PROGENy signatures across fibroblast populations. (C) Dot plot showing the pathway enrichment scores across fibroblasts populations and indications computed by pseudo-bulk FGSEA, for pathways with a p-value < 0.05. ECM, extracellular matrix; NES, normalized enrichment score; PROGENy, pathway responsive genes for activity inference; UMAP, Uniform Manifold Approximation and Projection; FGSEA, fast gene set enrichment analysis.

To validate these findings, we performed two complementary computational approaches. We first applied the algorithm CytoSPACE (21) to map to the cISH dataset two publicly available scRNAseq datasets of SSc and healthy skin (GSE195452 and GSE249279 (7, 8)). The ECM and collagens scores, as well as the TGFβ pathway activity computed from the enhanced cISH dataset were enriched in SFRP2+ RetD FB (Figure S9A), whereas the glycoproteins and proteoglycans scores were enriched in PI16+ FB (Figure S9B). We further performed spot deconvolution in the complementary spatial transcriptomic dataset, which confirmed that the SFRP2+ RetD and SFRP2+ PapD FB are in regions of collagens enrichment and of high TGFβ pathway activity (Figure S10).

We next performed functional enrichment analysis using the FGSEA and ULM methods via decoupleR (22). Both methods revealed that the fibroblast subsets have both shared and distinct biological functions (Figure 4C and Figure S11). The SFRP2+ RetD FB are profibrotic, ECM-producing fibroblasts. They demonstrated the highest enrichment scores for pathways related to fibrosis, such as “collagen formation”, “Response to TGFβ”, “Positive regulation of WNT signaling pathway”, and “IL-4 pathway” (Figure 4C and Figure S11A). These pathways were less enriched in the SFRP2+ PapD FB. In contrast, “Response to mechanical stimulus” was specifically enriched in SFRP2+ RetD FB (Figure S11A). SFRP2+ PapD FB demonstrated enriched pathways specific for this population, i.e. “signaling by PDGF”, “regulation of hemostasis” and de-enriched for “immune response” (Figure 4C and Figure S11A). CCL19+ nonPV FB are an inflammatory fibroblast population. Shared functions of CCL19+ PV and nonPV FB included several terms related to cytokine production and immune responses. Specific functions of CCL19+ PV FB included a high enrichment score on “regulation of monocyte chemotaxis” and de-enrichment of “immune system process” compared to the enrichment of “calcium ion homeostasis” and “lipid metabolic process” in CCL19+ nonPV FB (Figure 4C and Figure S11A). These differences highlight that the fibroblasts populations separated by BANKSY clustering are not only distinct with regards to their spatial location, but also with regards to their function.

The COL8A1+ FB may have broad spectrum of roles in modulating fibrotic and inflammatory responses as demonstrated enrichment in pathways related to collagen organization, but also in pathways such as “actin filament organization”, “regulation of lymphocyte migration” and “regulation of monocyte chemotaxis” (Figure 4C and Figure S11A). Furthermore, COL8A1+ FB demonstrated specific enrichment of “negative regulation of apoptotic signaling pathway”, suggesting resistance to apoptosis. The PI16+ FB demonstrated enrichment of “degradation of the extracellular matrix” and de-enrichment of “immune response”, suggesting homeostatic functions (Figure 4C and Figure S11A). Nevertheless, the PI16+ FB, the SFRP2+ PapD FB along with other fibroblast populations had higher enrichment scores for pathways related to inflammation and fibrosis in SSc, providing further evidence for functional reprogramming of individual fibroblast subpopulations in SSc (Figure S11B).

### Multicellular spatial domains in SSc skin

Given the profound effects of crosstalk with neighboring cells on cellular phenotype and function, we next aimed to characterize the local microenvironment of individual subpopulations of fibroblasts. We first annotated immune subpopulations by label transfer, with the scRNA-seq dataset GSE195452 as a reference. Visualization by UMAP revealed that the integration of the two datasets performed by scANVI effectively brought the immune cells from the cISH dataset in proximity to those from the scRNAseq in the latent space (Figure S12A). This enabled transfer of the labels from the reference dataset, with annotation of macrophages/monocytes, dendritic cells, mast cells, T cells, B cells, plasma cells and NK cells (Figure S12B).

We next aimed to identify multicellular spatial domains, or cellular neighborhoods (CNs), by clustering cells based on the composition of their neighboring cells. This approach yielded 14 CNs (Figure S13). We annotated the CNs according to their composition, i.e. to the cell population with the highest relative frequency inside each CN. Some of these CNs recovered histologically defined domains of the skin, e.g. the epithelium as Epithelial CN, or the vascular region as Endothelial CN (Figure S13). Most of the other CNs were mainly defined by the predominant fibroblast population, e.g. CCL19 nonPV CN, COL8 CN, SFRP2 CN, PI16 CN or CCL19 PV CN. SFRP2+ PapD, but not the SFRP2+ RetD FB were enriched in the Subepithelial CN, whereas the CCL19+ PV, but not the CCL19+ nonPV FB were enriched in the Endothelial CN, again confirming the spatial segregation of these populations with a distinct method (Figure S13).

The composition of several CNs were remarkably different in SSc compared to normal skin, with an enrichment of immune cell subsets in the CCL19 nonPV CN, COL8 CN and SFRP2 CN and with a de-enrichment of immune cells in the PI16 CN and CCL19 PV CN (Figure 5A). Furthermore, PI16+ FB were enriched in COL8 CN and SFRP2 CN in SSc, and thus in spatial proximity to SFRP2+ RetD and COL8A1+ FB, which indirectly supports the hypothesis that PI16+ FB can differentiate to pro-fibrotic fibroblast subsets in SSc (Figure 5A). These findings suggest profound changes in the spatial organization of SSc skin, where individual fibroblast subsets demonstrate not only aberrant changes in frequencies, but also in their spatial relationships with immune cells.

**Figure 5:**
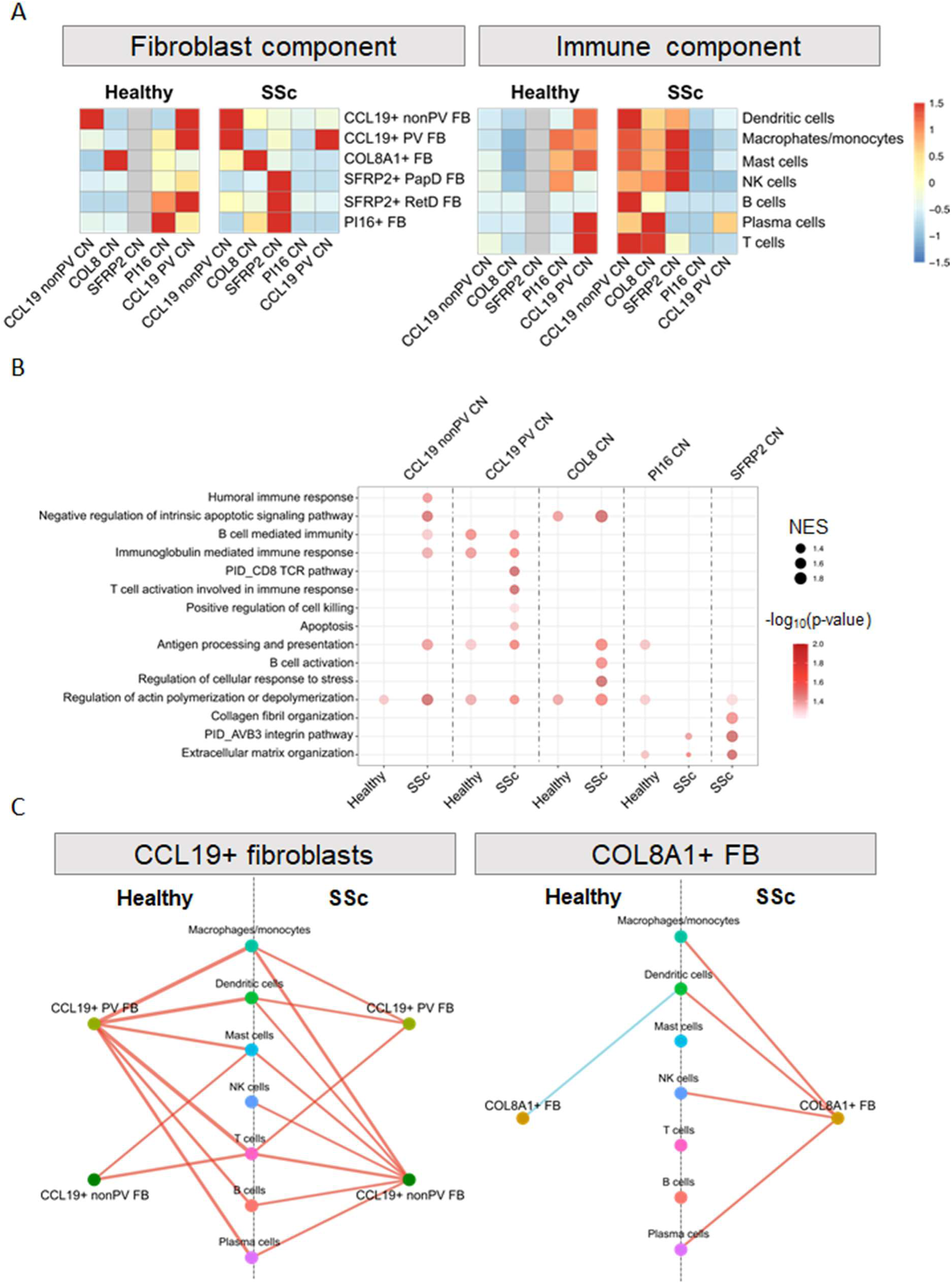
Changes in the cellular neighborhoods in SSc skin. (A) Heatmaps highlighting the shift in the cellular composition within the fibroblast CNs in healthy and SSc skin. The number of cells were normalized for each cell type across all CNs. SFRP2 CN was not detected in healthy samples and is shown in gray in the heatmap. (B) Dot plot showing the activation of immuno- and matrix remodeling pathways within fibroblast CNs in SSc compared to healthy controls. The activation of the pathways was analyzed by pseudo-bulk FGSEA. The size of the dot represents the NES, and the color represents the p-value. (C) Network visualization showing the immune interaction of indicated fibroblast populations in healthy and SSc skin by pairwise interaction analysis, based on the cellular distribution in the tissue. Cellular attraction (red) and avoidance (blue) are indicated in the plots. The width of the lines represents the proportion of samples exhibiting statistically significant interaction. The statistical significance was determined by FGSEA (B) or by permutation test (C). Results that did not reach statistical significance (p < 0.05) were excluded from the visualizations for clarity. SSc, systemic sclerosis; CN, cellular neighborhood; FGSEA, fast gene set enrichment analysis; NES, normalized enrichment score.

We reasoned that the changes in composition of CN in SSc can lead to functional changes within the local niches. Indeed, functional enrichment analysis demonstrated that fibroblast-enriched CNs, such as CCL19 nonPV CN, CCL19 PV CN, SFRP2 CN, PI16 CN or COL8 CN are functionally distinct in SSc compared to normal skin with changes in disease-relevant pathway enrichment scores (Figure 5B). Changes in SSc included the enhanced anti-apoptotic and antigen processing activities in CCL19 nonPV CN and COL8 CN in SSc. Activation of lymphocyte-mediated killing, including CD8 TCR signaling and apoptosis, was detected in CCL19+ PV CN in SSc. The SFRP2 CN was present exclusively in SSc skin and exhibited the highest scores in pathways related to ECM remodeling among the fibroblast-enriched CNs. These findings support the concept that the CNs identified are functional units within the skin that undergo pathologic changes in SSc.

### Cell-cell interaction and communication networks in SSc skin

We next aimed to unravel changes in cell-cell interaction networks in SSc skin. Pairwise interaction analysis revealed distinct cellular interaction networks in SSc skin (Figure 5C and Figure S14). In line with the findings on the immune composition in CNs, CCL19+ nonPV FB demonstrated increased interactions with several immune cell subsets, whereas CCL19 PV FB showed decreased the interaction with mast cells, B cells and plasma cells in SSc compared to normal skin (Figure 5C). COL8A1+ FB transitioned from an immune-inert state, as shown by the lack of interactions or even avoidance of immune cells, to an immune-engaging state, interacting with macrophages, dendritic cells, NK cells, and plasma cells in SSc (Figure 5C). In SSc skin, SFRP2+ PapD FB showed interactions with T cells and SFRP2+ RetD FB with mast cells and NK cells, all of which were not observed in normal skin (Figure S14).

We further performed spatially informed cell-cell communication analysis by CellChat (23), which infers interaction probabilities of multiple ligand-receptor (L-R) pairs, while considering the spatial proximity between cells. This analysis revealed profound changes in the patterns of L-R interactions in SSc compared to control skin (Figure 6, Figure S15 and Figure S16). SFRP2+ RetD FB demonstrated L-R interactions with several other fibroblast subsets and with mast cells, NK cells, or dendritic cells in SSc, whereas in control skin they were only communicating with other SFRP2+ RetD FB (Figure 6A). In contrast, the SFRP2+ PapD FB lost communication with multiple cell types in SSc (Figure 6A). Of note, CellChat analysis without spatial distance constraints, which did not account for spatial information, did not resolve these differences between SSc and normal skin and showed comparable L-R interactions of both SFRP2+ FB subsets (Figure S15A).

**Figure 6:**
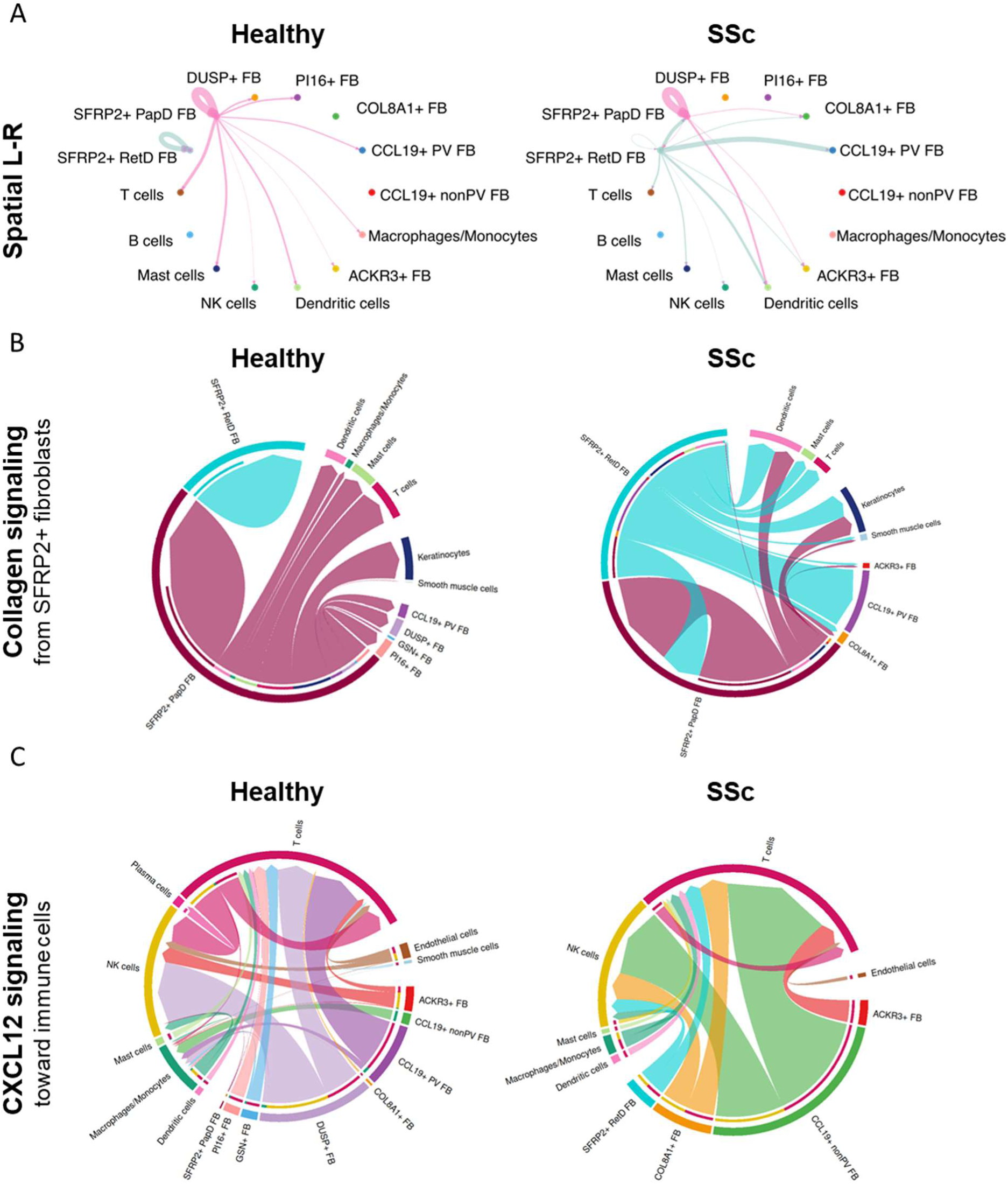
Distinct cell-cell communication in SSc identified by spatially informed ligand-receptor analysis. (A) Representative circle plots showing the interaction of SFRP2-expressing fibroblasts in healthy and SSc skin by spatial ligand-receptor analysis using CellChat. The distance constraints were set to 250 µm for secreted signaling and 20 µm for contact-dependent signaling. Interaction routes are represented by lines, with line width indicating interaction strength as computed by CellChat. (B) Chord diagrams showing the collagen signaling from the SFRP2-expressing fibroblasts. (C) Chord diagrams showing the CXCL12 signaling received by immune cells and their source. In chord diagrams (B and C), the thinner inner bars represent the signaling strength received by the indicated targets. SSc, systemic sclerosis; L-R, ligand-receptor analysis.

At the signaling level, SFRP2+ RetD FB in SSc demonstrated an increased relative strength for outgoing, but a decreased strength for incoming signals for COLLAGEN and FN1 compared to normal skin, which may point towards a reduced capacity for ECM sensing in SSc (Figure 6B, S15B). Indeed, further analysis indicated decreased ECM signals received by CD44, a receptor with well-characterized roles in ECM sensing (24), on SFRP2+ RetD FB compared to other fibroblast populations in SSc (Figure S16A). Furthermore, the CCL19+ nonPV FB demonstrated an increased relative strength for outgoing signals for CXCL12 in SSc, suggesting a higher chemotactic capacity (Figure 5C and S16C). PDGF signaling was altered in SSc skin: fibroblast subsets with increased frequency in SSc, such as SFRP2+ RetD and COL8A1+ FB started to receive PDGF signals in SSc, whereas in control skin only fibroblast subsets with lower frequency in SSc, such as SFRP2+ PapD and CCL19+ PV were receiving PDGF signals, indicating a switch from homeostatic to pro-fibrotic roles of PDGF signaling in SSc (Figure S15D). Furthermore, PDGF signals were sent by endothelial cells in SSc and by monocytes/macrophages in control skin.

These spatially resolved analyses demonstrate major changes in signaling between distinct fibroblast subsets and other cells in SSc skin that could not be resolved in non-spatial datasets.

### Predictors of progression of skin fibrosis in SSc

We investigated whether the frequencies of fibroblast subpopulations - either on their own or stratified by their interaction partners - at the time of biopsy could predict the progression of skin fibrosis at follow-up. Patients with progression of skin fibrosis at follow-up had a higher frequency of COL8A1+ FB than those with stable skin fibrosis (Figure 7A). Other fibroblast populations were not significantly different between the two groups (Figure S17). Furthermore, COL8A1+ FB in close spatial proximity to macrophages/monocytes, mast cells or B cells had a significantly higher frequency in patients with progression of skin fibrosis, whereas COL8A1+ FB with spatial proximity to other immune cell populations demonstrated only milder differences between the two groups (Figure 7B-D and Figure S18A-D).

**Figure 7:**
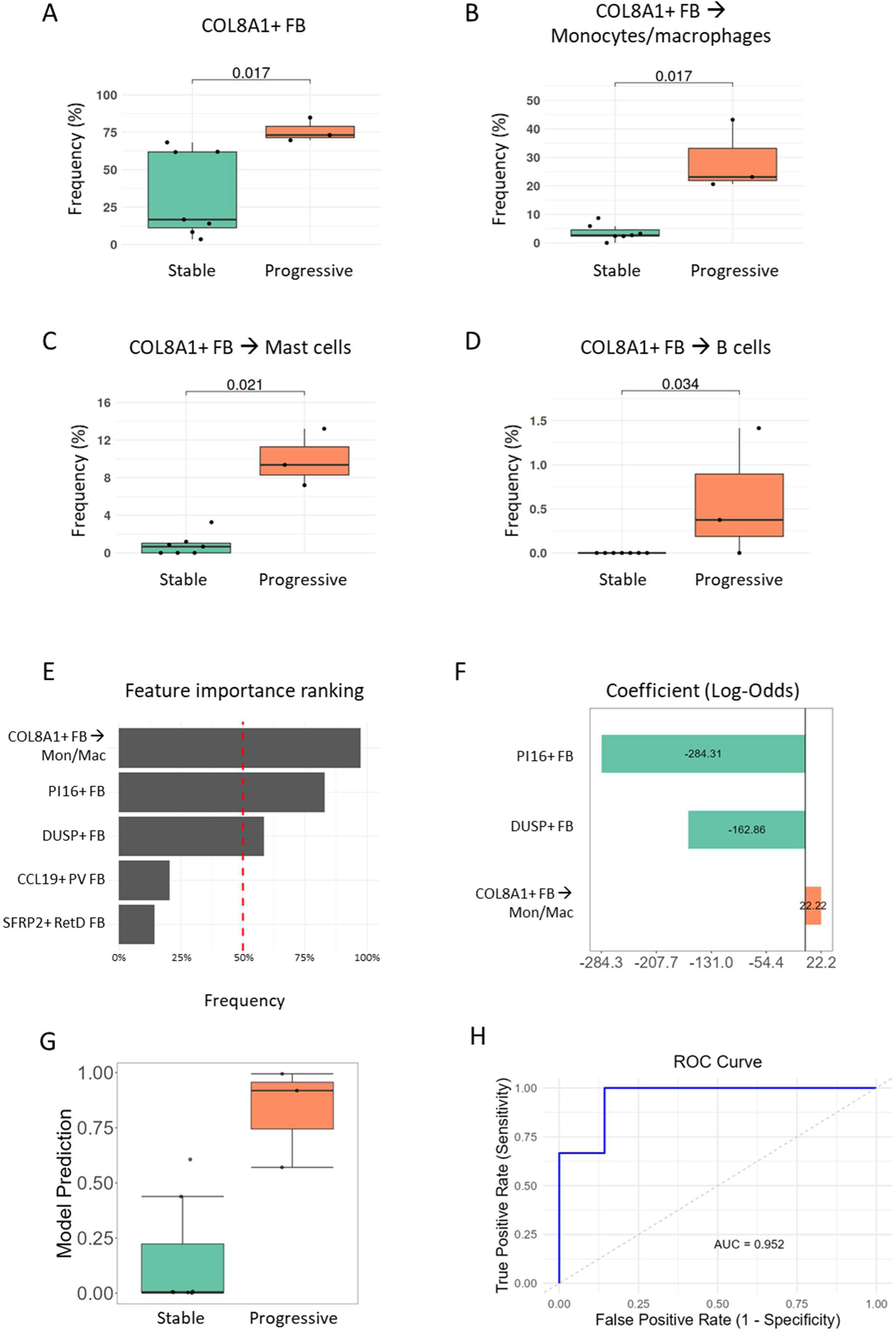
Predictors of progression of skin fibrosis in SSc. (A-D) Frequencies of all COL8A1+ FB (A) and of COL8A1+ FB in spatial proximity to monocytes/macrophages (B), to mast cells (C) or to B cells (D) among all dermal cells. (E) Top highly ranked features predictive of progression of skin fibrosis after bootstrapping with LASSO regression, starting from the frequencies of COL8A1+ FB in spatial proximity to monocytes/macrophages or to mast cells or B cells, along with the frequencies of SFRP2+ RetD FB, of PI16+ FB, of DUSP+ FB and of CCL19+ PV FB as predictors. (F) Increased frequencies of COL8A1+ fibroblasts in close spatial proximity to monocytes/macrophages were associated with a higher likelihood of progressive skin fibrosis, whereas elevated frequencies of PI16+ FB and DUSP+ FB were linked to a decreased likelihood of fibrosis progression. (G) Model prediction results for each donor. (H) ROC curve and its AUC for evaluation of model performance. FB, fibroblasts; RetD, reticular dermis; LASSO, least absolute shrinkage and selection operator; ROC, receiver operating characteristic; AUC, area under the curve.

We further generated two LASSO predictive regression models with progression of skin fibrosis at follow-up as outcome. The first model that included the frequencies of the fibroblast subsets with shifts in frequencies in SSc skin as predictors for progression of skin fibrosis performed with a moderate accuracy, with a significant overlap between the predicted stable and progressive patients, and with an AUC of 81% (Figure S19A-D). In a second model, we included the frequencies of COL8A1+ FB with spatial proximity to either macrophages/monocytes, or to mast cells or to B cells instead of the frequencies of all COL8A1+ FB, while keeping the other predictors unchanged. This model demonstrated improved prediction accuracy, allowing a better separation between the predicted progressors and predicted stable patients, with an AUC of 95% (Figure 7E-H). The frequency of COL8A1+ in proximity to monocytes/macrophages was the most important feature of the second model and increases in their frequency were predictors of progression, whereas the frequency of PI16+ fibroblasts were the second most important feature and increases in their frequency were associated with stable skin fibrosis (Figure 7E, F). This suggests an active differentiation through the trajectory starting from PI16+ FB and ending on COL8A1+ FB at early stages during progression of skin fibrosis. These findings highlight that incorporation of spatial information can improve the prediction of skin fibrosis compared to analyzing changes in cell frequencies alone.

## Discussion

We present herein a novel approach for identification and functional characterization of cell populations based on transcriptional profiles as well as on spatial localization using cISH as an emerging technique, in conjunction with novel bioinformatical pipelines. We demonstrate the validity of this approach by confirming deregulation of previously identified subpopulations of fibroblasts. Moreover, we complement the cISH-based approach with analysis of a spatial transcriptomics dataset from a second cohort of SSc patients and controls and with integration with publicly available scRNA-seq datasets from further SSc cohorts to combine the strengths of these different approaches. The cISH-based approach combines high, subcellular resolution of cISH with transcriptional phenotyping. The spatial RNA sequencing approach by Visium enables a deeper, transcriptome-wide phenotyping, but at a lower spatial resolution, where one spot corresponds to approximately four cells on average. Moreover, the cISH and Visium datasets were also complementary with regards to the patient cohorts – the cISH dataset included Caucasian patients, whereas the spatial transcriptomics dataset included Han Chinese patients.

Analyses of our cISH data using the BANKSY algorithm, which considers expression levels as well as spatial information, enabled us to identify nine different fibroblast subpopulations in SSc and control skin. Identification of well-known fibroblast populations such as PI16+, SFRP2+ or CCL19+ fibroblasts validated our approach (6, 7, 9). Our BANKSY-based spatial phenotyping approach further segregated SFRP2+ FB and CCL19+ FB into four distinct populations based on the phenotypes in the local microenvironment. SFRP2+ FB consisted of an SFRP2+ RetD population with localization in the reticular layers of the dermis and a population of SFRP2+ PapD with cells being localized only in the upper dermis. CCL19+ FB consisted of a CCL19+ PV subpopulation with perivascular localization and another population of CCL19+ nonPV, which is diffusely distributed within the dermis. We validated the distinct spatial localization of these populations by two complementary methods, i.e. by spot deconvolution of the Visium dataset and by the neighbourhood analysis.

Of note, these spatially separated fibroblast populations do not only show differences in spatial localization, but are also functionally distinct with enrichment of distinct functional terms. For example, in SFRP2+ PapD FB regulation of hemostasis was specifically enriched, whereas in SFRP2+ RetD FB the response to mechanical stimulus was specifically enriched, and TGFβ signaling was stronger than in SFRP2+ PapD FB. The two populations of CCL19+ FB also showed profound functional differences. Moreover, these pairs of fibroblasts segregated by spatially informed phenotyping were also localized in distinct cellular neighborhoods and differed with regards to cellular interactions. SFRP2+ RetD FB interacted with a wide range of immune cells. On the other hand, only a small proportion of dendritic cells were found in the surroundings of SFRP2+ PapD FB. Finally, these populations also displayed clear differences in ligand-receptor interactions. E.g., ECM sensing by CD44 was specifically decreased in SFRP2+ RetD in SSc; furthermore, only CCL19+ nonPV, but not CCL19+ PV, signaled to NK cells via CXCL12-ACKR3. The novel, spatially defined populations of SFRP2+ RetD and SFRP2+ PapD as well as CCL19+ PV and CCL19+ nonPV fibroblasts are thus functionally clearly distinct with differences in unique gene expression patterns, signaling pathways and cellular interactions. Furthermore, these populations also show specific alterations in SSc skin, providing further evidence for distinct roles in the pathogenesis of SSc.

Our study also provides clear evidence that the profibrotic phenotype of fibroblasts in SSc is the result of the combined effects of different processes: 1.) Our study demonstrates changes in the frequencies of fibroblast populations with increased frequencies in profibrotic populations. These changes include the elevated abundance of SFRP2+ RetD FB, which had demonstrated the highest collagen expression in our dataset. Shift toward profibrotic subsets have also been reported in non-spatially resolved scRNA-seq studies in SSc (6–10). The changes in frequencies might be partially caused by a shift in the differentiation trajectory of precursor fibroblasts, i.p. of PI16+ FB, towards pro-fibrotic subsets. 2.) In addition, we demonstrate that several individual populations are further biased towards a more profibrotic phenotype in SSc skin with upregulation of profibrotic genes and pathways, including TGFβ, WNT, and ECM-modulating pathways. 3.) The local neighborhoods change drastically in SSc, thereby providing a microenvironment that fosters development of profibrotic fibroblasts. We detected upregulation of immune cell subsets with potential pro-fibrotic roles such as monocytes/macrophages, mast cells or plasma cells in the CCL19+ nonPV, COL8, and SFRP2 CN in SSc. These changes in the interactions with immune cells are not a simple reflection of increased leukocyte counts in SSc skin, as mentioned interactions are specific for certain leukocyte subsets and occur only with certain fibroblast populations. In contrast, other fibroblast subsets such as CCL19+ PV FB demonstrate decreased interactions with immune cells in SSc.

The cISH with high spatial resolution enables studies of alterations in signaling pathways within local niches in SSc and analyses of heterogeneity of pathway activation across different multicellular spatial domains. We provide first evidence for specific alterations of disease-relevant signaling pathways within local tissue niches that accompany the changes in composition of niches in SSc skin: the COL8 CN and the CCL19 nonPV CN had enhanced anti-apoptotic and antigen processing activities, whereas the CCL19+ PV CN demonstrated activation of CD8 TCR signaling and cell killing in SSc. Furthermore, the SFRP2 CN, which is present exclusively in SSc skin, demonstrated profound enrichment of pathways related to ECM remodeling in comparison to other fibroblast-enriched niches.

We also provide first evidence that the frequencies of two fibroblast populations and their interactions with immune cells may help to predict the future course of skin fibrosis. The frequencies of COL8A1+ FB were associated with future progression of skin fibrosis, whereas PI16+ FB were associated with stable disease. This correlates well with the functional role of these two fibroblast subpopulations and their developmental trajectory. COL8A1+ FB demonstrate enrichment in numerous pathways related to both fibrosis and inflammation.

COL8A1+ FB are also the end stage of differentiation of PI16+ FB through a trajectory that involved SFRP2+ RetD FB, another population with altered frequencies in SSc. This suggests that the active differentiation of PI16+ FB towards COL8A1+ FB, with potential recruitment of immune cells by the COL8A1+ FB, might drive progression of skin fibrosis. In line with this, the frequency of COL8A1+ interacting with monocytes/macrophages provided an improved predictive power compared to the frequencies of all COL8A1+ fibroblasts alone. Thus, by providing spatial information, our approach could provide an improved risk stratification than that based on evaluation of cell frequencies alone.

In summary, we used cISH in conjunction with novel bioinformatical pipelines that consider spatial context to identify four novel fibroblast subpopulations and to study the microenvironment and cellular interactions of fibroblasts in SSc and normal skin. We characterize SFRP2+ RetD FB as prototypical profibrotic, ECM producing population that resides within a specific inflammatory niche, CCL19+ nonPV FBs as pro-inflammatory population that attracts and activates immune cells and COL8A1+ FB as a population with roles in both ECM remodeling as well as immune cell recruitment that is associated with progression of skin fibrosis. These results may provide a rationale for specific therapeutic targeting of SFRP2+ RetD-, CCL19+ nonPV- and COL8A1+ fibroblasts as disease-promoting fibroblast subsets and may stimulate follow-up studies to confirm the potential of COL8A1+ FB counts as a marker for fibrosis progression.

## Methods

### Study subjects and clinical data

All twelve SSc subjects fulfilled the American College of Rheumatology (ACR) / European Alliance of Associations for Rheumatology (EULAR) 2013 criteria (25). Skin biopsies of eight healthy donors matched for age and gender served as controls. All SSc patients and controls provided signed informed consent forms approved by the ethical committee of the Medical Faculties of the University Hospitals of Düsseldorf, Erlangen, Cologne or Tübingen Universities. All skin biopsies of patients with SSc were obtained from the upper limb, at least 2 cm proximal from the styloid process of the ulna, using a disposable biopsy punch. Skin biopsies from healthy donors were collected from patients undergoing surgery for either trauma (without any injury at the region where the biopsy was taken from) or degenerative joint disease (osteoarthritis).

Early SSc was characterized as a disease duration of less than three years from the onset of the first non-Raynaud phenomenon symptom. Disease activity was determined using the EUSTAR disease activity index. A cut-off value of ≥2.5 is used to define a patient as having active disease (26). The inflammatory phenotype was defined by the presence of CRP levels > 6 mg/l, elevated thrombocyte counts over 330,000/µL, arthritis, or myositis (27). mRSS evaluations were conducted by experienced and trained physicians. ILD progression was assessed based on the criteria established in the INBUILD trial (28). Progressive skin fibrosis was defined as at least a 25% increase in mRSS from 6 months before biopsy to baseline, as well as measured during a follow-up visit conducted 6 months after biopsy. Demographic and clinical characteristics of the SSc patients are summarized in Table S1.

### Sex as a biological variable

We examined the samples obtained from both male and female donors. The detailed demographic data of the donors is described in **Table S1**.

### Sample preparation

Formalin-fixed paraffin-embedded (FFPE) 5μm skin sections from the human biopsies were prepared. Consecutive sections were used to generate a hematoxylin and eosin (H&E) staining for evaluation of histopathologic changes, as well as to select the region of interest to be analyzed by cISH.

### Acquisition of cISH

To investigate the transcriptional changes with spatial context at single-cell resolution, we applied imaging-based spatial transcriptomics on the FFPE skin samples. In brief, cISH was performed by NanoString Technologies (now Bruker Spatial Biology, Inc) using CosMx SMI platform as described (29). An mRNA probe panel targeting 969 genes was created by combining CosMx Universal Cell Characterization RNA Core Panel with 86 custom-designed probes targeting 19 marker genes of fibroblast populations in SSc skin (Table S2). Negative probes targeting 20 negative control sequences obtained from External RNA Controls Consortium (ERCC) were also included in the panel as previously described (30). Marker genes for fibroblast populations were selected based on publicly available scRNA-seq SSc skin datasets using COMET and sc2marker (6, 8, 31, 32). Cell segmentation was performed on the CosMx SMI platform, which uses a pretrained model based on Cellpose algorithm utilizing cellular membrane, nuclear, and RNA staining (30). Transcript locations and single-cell expression matrices were obtained from the AtoMx SIP platform using the Export and Flat File Export modules.

### cISH analysis

#### Quality control and data normalization

The flat files from AtoMx were imported as a SpatialFeatureExperiment object using Voyager R package (33). Poorly segmented objected were excluded based on DAPI staining intensity and object sizes. Cells with low RNA counts (≤ 6) and high proportion of negative control counts (≥ 10%) were excluded for the further analysis. Expression values were normalized by dividing each cell’s raw RNA counts by its total counts, with a lower threshold of 20 to prevent excessive upscaling of low-count cells following the instruction from Brucker Spatial Biology (https://nanostring-biostats.github.io/CosMx-Analysis-Scratch-Space/). A z-score scaled count matrix was obtained by performing gene-wise normalization on the normalized expression data.

#### Spatially informed cell phenotyping

To identify populations using both gene expression profile and spatial information, we performed spatial clustering using the Banksy R package (11). In brief, we constructed a Building Aggregates with a Neighborhood Kernel and Spatial Yardstick (BANKSY) expression matrix that incorporated both the self-expression profile of each cell and the expression profiles of its neighbors before performing dimensionality reduction. The neighbor expression profiles were derived using two metrics: a weighted mean expression and a gradient expression.

The mean expression was weighted based on the distances between the cells. The weight decreased exponentially with distances and was normalized by the median distance among the neighbors. The gradient expression was computed similarly to the mean expression but includes an additional azimuthal component which accounted for the directional information from the neighbors. The neighboring cells were defined by k-nearest neighbors (kNN) with k = 15 for the mean expression and k = 30 for the gradient expression.

A neighbor-augmented expression was then generated by combining the cells’ own expression and two neighbor expression matrices with the spatial mixing factor λ = 0.25. Nonspatial clustering was also performed by setting λ = 0 for comparison. To identify the cell populations, we further performed dimensionality reduction by principal component analysis (PCA) and Leiden clustering with the first 20 PCs. Uniform Manifold Approximation and Projections (UMAP) were created from the first 20 PCs for the data visualization.

#### Pseudotime trajectory inference

Pseudotime trajectories for the fibroblast differentiation were constructed using the R package Monocle (v3) (15, 34, 35). For pseudotime construction, the same assay and reduced dimensions were used as for BANKSY clustering. The orders of the cells along pseudotime were determined by the “*orderCells*” function, with PI16+ FB as the root.

#### Cell phenotyping by label transfer

Annotation of immune populations was conducted by label transfer with publicly available SSc skin scRNA-seq dataset (GEO accession GSE195452) using the popV python package (36). The batch balanced KNN (BBKNN), Scanorama, and random forest (RF) classifier showed limited convergence of the cells across two datasets in the integrated latent space. Decent integration was found in the rest of the methods, including support vector machine (SVM) classifier, single-cell ANnotation using Variational Inference (scANVI), and kNN algorithm on harmony- or single-cell variational inference-(scVI) integrated data. Among these methods, scANVI yielded the most robust results for label transfer on our CosMx dataset. We further validated the label transfer approach by using Leiden clustering, with cluster annotations based on marker genes closely aligning with scANVI-predicted labels.

#### Functional analysis

Normalized count data was used for expression profiling. Marker gene expression for each cell population was visualized using heatmaps and UMAP plots. The marker genes were discovered by Wilcoxon rank sum tests implemented in the Seurat R package with the “*FindAllMarkers*” function (37). To address the sparsity inherent to spatial transcriptomics datasets, smoothed expression values were calculated using kNN in the UMAP space, following the instruction from Bruker Spatial Biology, Inc. (https://nanostring-biostats.github.io/CosMx-Analysis-Scratch-Space/) for the visualization of gene expression on UMAP plots.

To characterize fibroblast populations, we computed expression scores for extracellular matrix (ECM) components, including collagen, using the “*AddModuleScore*” function in the Seurat R package. The ECM geneset used for the scores was obtained from the Core Matrisome Annotations (19). To explore the biological functions across populations and cellular neighborhoods, we generated pseudo-bulk expression profiles by averaging expression within defined cell populations or conditions. The pathway activity scores were then computed by Fast Gene Set Enrichment Analysis (FGSEA) and univariate linear model (ULM) using decoupleR R package, with a curated gene set obtained from Bader Lab (version Human_GOBP_AllPathways_noPFOCR_no_GO_iea_November_04_2024) (22, 38, 39). TGFβ activity score was inferred using a multivariate linear model with PROGENy gene signatures (20). The top 2000 gene weights were retrieved from PROGENy using the OmnipathR R package (40). The gene sets and weights were filtered to match the genes included in the CosMx panel for the functional analysis.

To investigate cell-cell communication, we conducted ligand-receptor analysis using the CellChat R package (23). Ligand-receptor interaction data was sourced from CellChatDB database. We performed the analysis in both spatial and non-spatial scRNA-seq-like modes to evaluate the added insights provided by spatial information. For both types of analysis, the averaged expression was computed by truncated mean, with trimming proportion set to 10%. In the spatial mode of the analysis, we set distance constraints of 250 µm for secreted signaling and 20 µm for cell contact-dependent signaling. Interactions involving fewer than five cells were excluded from further analysis to ensure robust communication detection.

### Cellular neighborhoods and interaction analysis

Cellular neighbors were defined as adjacent cells identified through Delaunay triangulation, with centroid distances within 50 µm. Neighbour cell pairs were detected using the imcRtools R package (41). To investigate the cellular local environments, we identified cellular neighborhoods (CNs) as previously described (14, 42). Briefly, we first computed the composition of neighboring cells for each cell, and K-means clustering was applied to these compositions to define CNs. The cells residing in the same CN shared similar neighboring cell types. We created heatmaps to visualize the cell composition within CNs. The heatmaps were scaled according to the proportion of individual cell type in the entire dataset, unless otherwise specified.

To analyze the spatial proximity between cells, we performed cellular pairwise interaction analysis using the imcRtools R package. The significance of individual pairwise interactions was determined using permutation testing. These tests calculated the number of neighboring cells for each subpopulation and compared it to an empirical null distribution obtained by randomly shuffling cell type labels for 1000 permutations. A pairwise interaction with a *p*-value below 0.05 was considered as significant. We interpret statistically significant higher observed cell counts as cellular attraction and lower counts as cellular avoidance. Dot plots and interaction network visualizations were generated using ggplot2 and ggraph R packages.

### Multimodal integration

#### scRNA-seq analysis

We retrieved publicly available scRNA-seq datasets for SSc skin (GEO accession GSE195452, GSE138669, GSE249279) (6–9). The cells were annotated based on the provided cell labels from the source datasets (GSE195452 and GSE138669) or based on the marker genes provided in the original publication (GSE249279). Raw count data was used for PopV-based label transfer and integration as described above. To validate the phenotype of fibroblast populations detected by CosMx, we conducted CytoSPACE-based integration between CosMx and scRNAseq datasets (21). CytoSPACE was run in single-cell mode, using coarse cell labels prior information to account for variations in the cell type composition across the datasets.

#### Spatial sequencing analysis

Sequencing-based spatial transcriptomics was performed on the skin FFPE sections obtained from donors at Huashan Hospital by Visium platform. To map the cell populations and cellular neighborhoods identified in CosMx onto Visium dataset, we performed anchor-based reference mapping using Seurat R package. This approach enables probabilistic label transfer, accommodating multiple cells per spot in the Visium dataset. Both datasets were normalized using the *SCTransform* function, with the clip range set to [-10, 10] for the CosMx dataset following the recommendations of Seurat. The labels were transferred from CosMx dataset to Visium dataset based on the anchors computed from the expression of 909 shared genes between two datasets.

#### Predictive modeling

LASSO (least absolute shrinkage and selection operator) regression was used to build two models that predict progression of skin fibrosis at a follow-up of six months after biopsy, as described (43). Ten-fold cross-validation was used to find the optimal λ value (regularization coefficient). A first model was built with the frequencies of fibroblasts found to be statistically significant in the pairwise comparisons as predictors (COL8A1+, SFRP2+ RetD, PI16+, DUSP+ and CCL19 PV FB). A second model was built with the same predictors as the first model but replacing the frequencies of all COL8A1+ FB with the frequencies of COL8A1+ FB that are in spatial proximity with either monocytes/macrophages, or with mast cells or B cells. Bootstrapping (sampling with replacement, 1000 iterations) was used to rank the importance of all features. Variables with ≥50% frequency were selected for training in the final model. Data were randomly split into training and validation sets at a 75:25 ratio (1000 iterations) for cross validation and evaluation of model performance. In each iteration, progression of skin fibrosis was predicted on the validation set. Receiver operating characteristic (ROC) curve was calculated from the median prediction of each patient.

## Statistics

Data are presented as dot plots with the median ± interquartile range (IQR). Each dot represents one biological replicate, unless indicated otherwise in the figure legend. Protein expression levels are shown as heatmaps representing the mean of the z-score-normalized values. GraphPad Prism 8 was used to generate dot plots and to perform statistical analyses. Heatmaps were generated using the imcRtools R package (41). Mann-Whitney U-tests were applied for the comparison between two groups, if not indicated otherwise.

## Study approval

This study has been approved by the Ethics Committee of the Faculty of Medicine at Heinrich Heine University Düsseldorf (approval number: 2022-2189_3, 2023-2561).

## Data availability

Data are available upon reasonable request to the corresponding authors.

## Author contributions

YNL, JHWD and AEM designed the study. YNL, TF, AHG, ML, VD, AM, HC, and AEM were involved in acquisition and analysis of data. YNL, TF, AHG, ML, CB, ANP, TK, JH, AK, GS, SD, JHWD and AEM were involved in interpretation of data. ML, CB, ACP, TK, JH, AK, and GS provided essential samples. All authors were involved in manuscript preparation and proof-reading.

## Acknowledgments

We thank Christoph Liebel, Philipp Steinbrecher and Lukas Sokolowski for excellent technical assistance. Michael Bailey, Trieu My Van, and Michael Leon from NanoString Technologies (now Bruker Spatial Biology) for supporting the CosMx runs. The project was supported by the following grants: Grants DI 1537/17-1, DI 1537/20-1, DI 1537/22-1, DI 1537/23-1 of the German Research Foundation, an unrestricted research grant from the Hiller-Foundation (JHWD), MA 9219/2-1 of the German Research Foundation (AEM), grants 2021_EKEA.03 (AEM) and 2022_EKMS.02 (AEM) of the Else-Kröner-Fresenius-Foundation, The Edith Busch and World Scleroderma Foundation Research Grant Programme 2022-2023 (AEM) and the Research Committee of the Medical Faculty of the Heinrich-Heine University Düsseldorf (Forschungskommission; ID 2022-18, ID 2023-33 and ID 2023-31 to AHG, AEM and JHWD, respectively). Our analysis was supported by a de.NBI Cloud project (YNL) within the German Network for Bioinformatics Infrastructure (de.NBI) and ELIXIR-DE (Forschungszentrum Jülich and W-de.NBI-001, W-de.NBI-004, W-de.NBI-008, W-de.NBI-010, W-de.NBI-013, W-de.NBI-014, W-de.NBI-016, W-de.NBI-022).

## Supplementary figures

**Figure S1:**
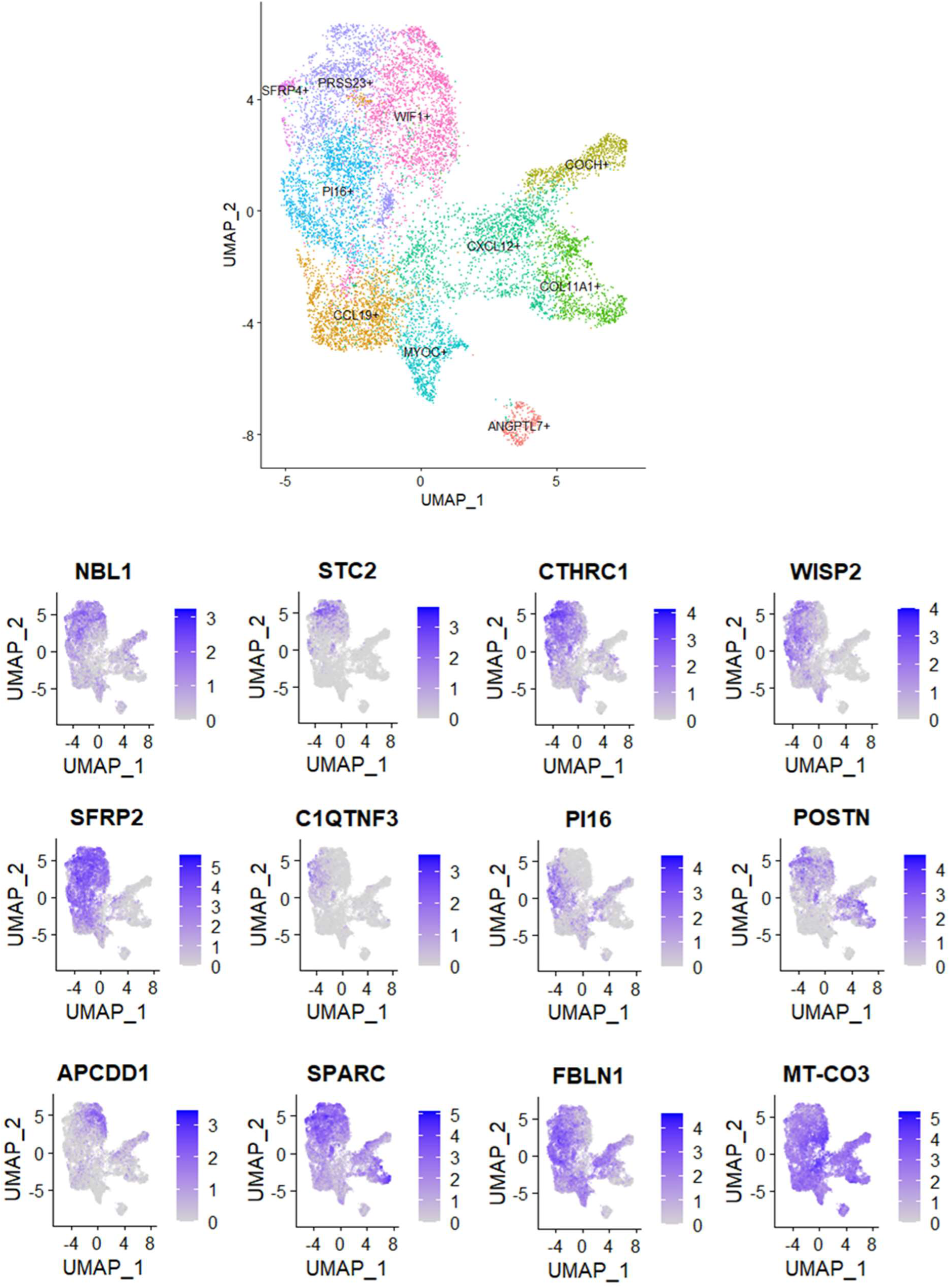
Marker gene selection from publicly available scRNA-Seq SSc skin dataset for cISH assay. UMAP plots demonstrating the expression of selected add-on genes to better characterize fibroblast populations in SSc skin. The fibroblast populations identified by our previous analysis on the scRNA-seq dataset (accession number GSE138699) are shown in the upper panel. The expression of selected marker genes is shown in the lower panel. UMAP, Uniform Manifold Approximation and Projection.

**Figure S2:**
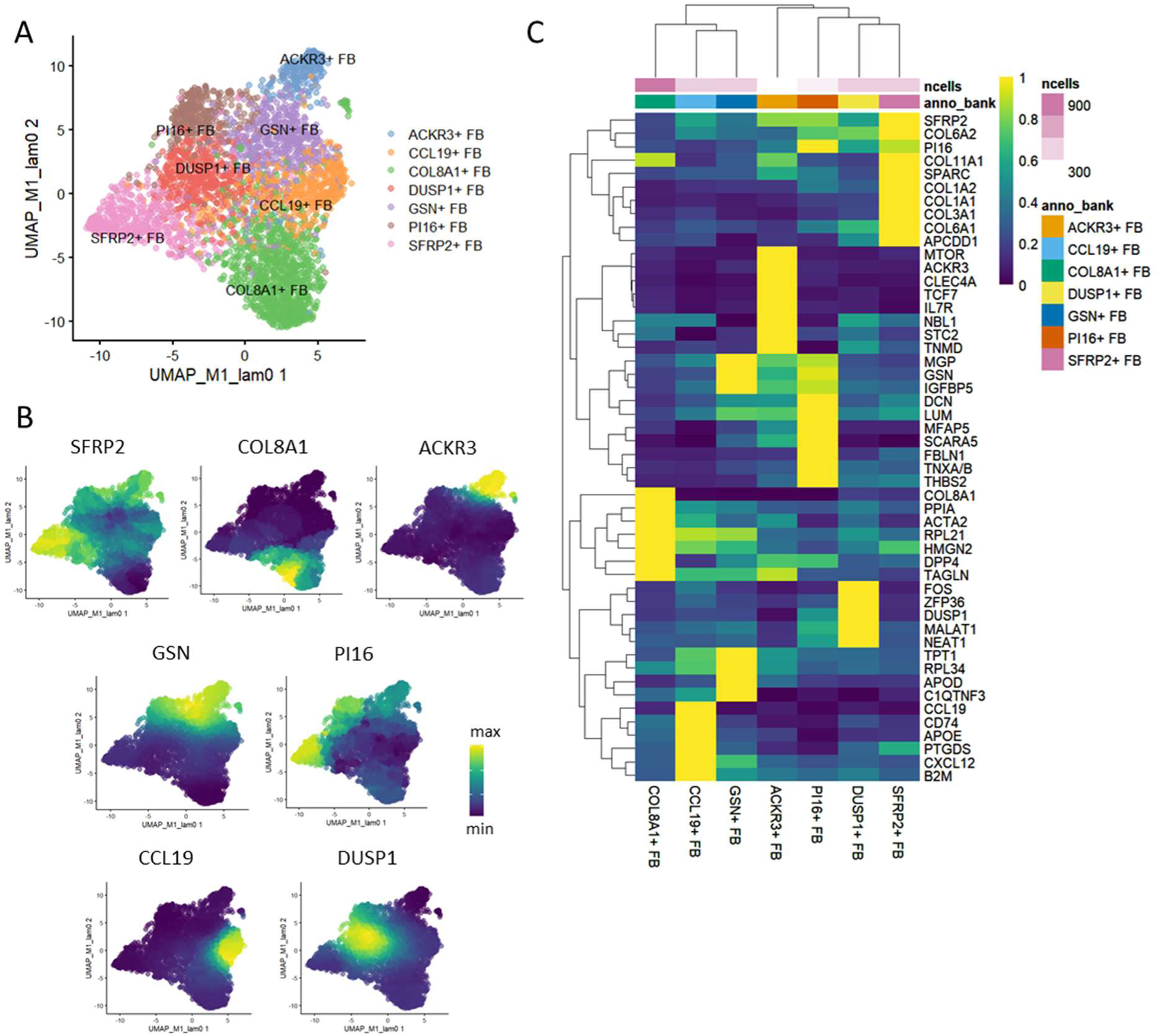
Fibroblasts population identified by non-spatial phenotyping. (A) UMAP plot showing the fibroblast populations identified by conventional non-spatial Leiden clustering in skin tissue. (B, C) The expression of marker genes of non-spatial fibroblast clusters is shown in UMAP plots and expression heatmap (C). The expression in UMAP was smoothed by the neighboring cells in the latent spaces. UMAP, Uniform Manifold Approximation and Projection.

**Figure S3:**
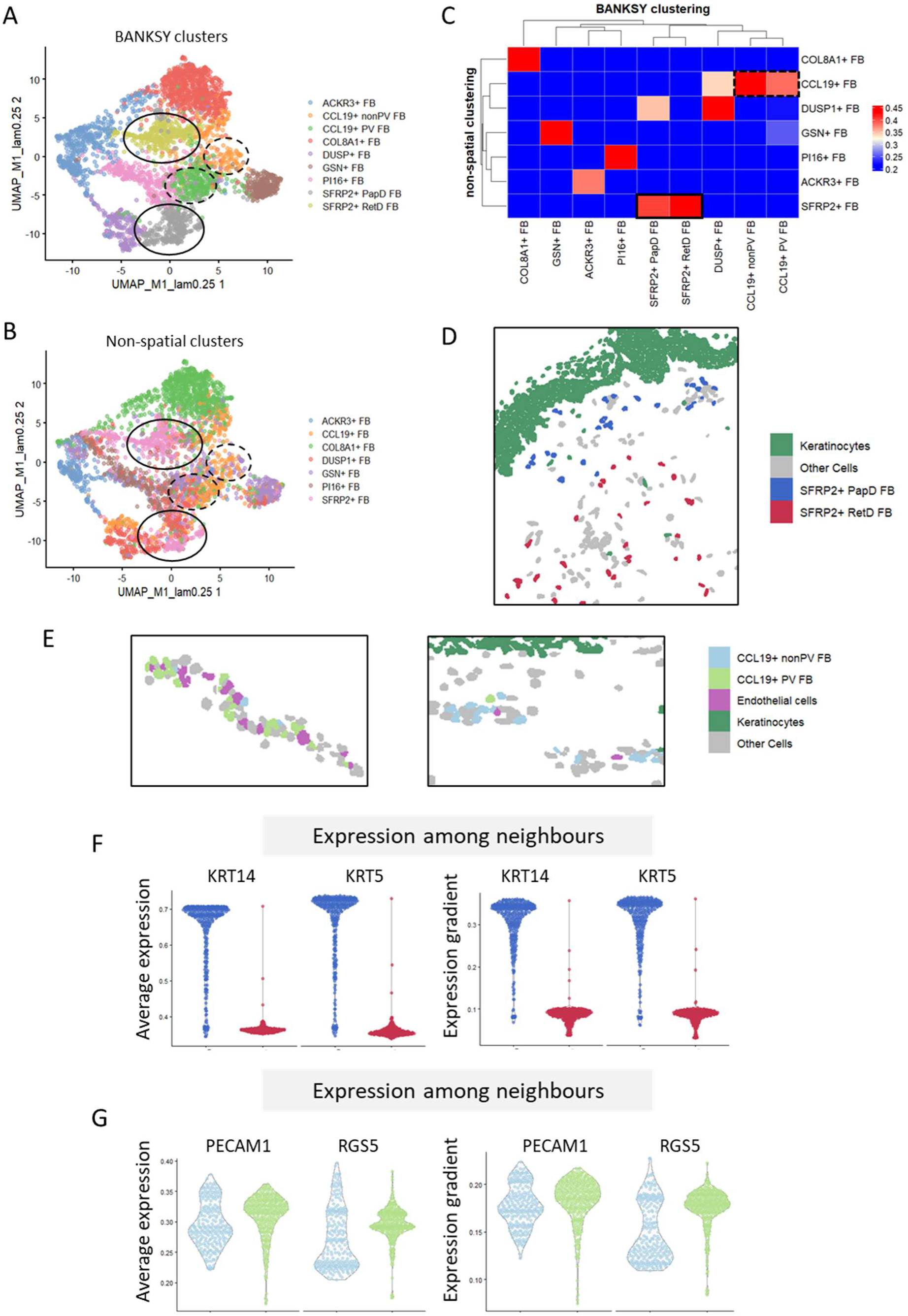
Characterization of fibroblast populations identified by non-spatial and spatial clustering. (A, B) UMAP plots showing the fibroblast populations identified by spatial BANKSY (A) and non-spatial (B) clustering. Both UMAP plots were generated using BANKSY expression matrix for comparison. The solid circles highlight the correspondence of SFRP2+ FB in non-spatial clustering to two clusters from BANKSY spatial clustering, SFRP2+ PapD FB and SFRP2+ RetD FB. The correspondence of CCL19+ FB from non-spatial clustering to CCL19+ PV FB and CCL19+ nonPV FB obtained from spatial clustering was highlighted by dashed circles in the UMAP plots. (C) A heatmap showing the correspondence of fibroblast populations identified by non-spatial and spatial clustering. (D, E) Representative images of the distribution of the fibroblast populations that were segregated by BANKSY clustering and their surroundings. (F, G) The averaged expression and the expression gradient of epithelial markers *KRT14* and *KRT5* (F), and vascular markers *PECAM1* and *RGS5* (G) among the neighbors computed by BANKSY on SFRP2+ BANKSY clusters (F) and CCL19+ BANKSY clusters (G). UMAP, Uniform Manifold Approximation and Projection; BANKSY, Building Aggregates with a Neighborhood Kernel and Spatial Yardstick.

**Figure S4:**
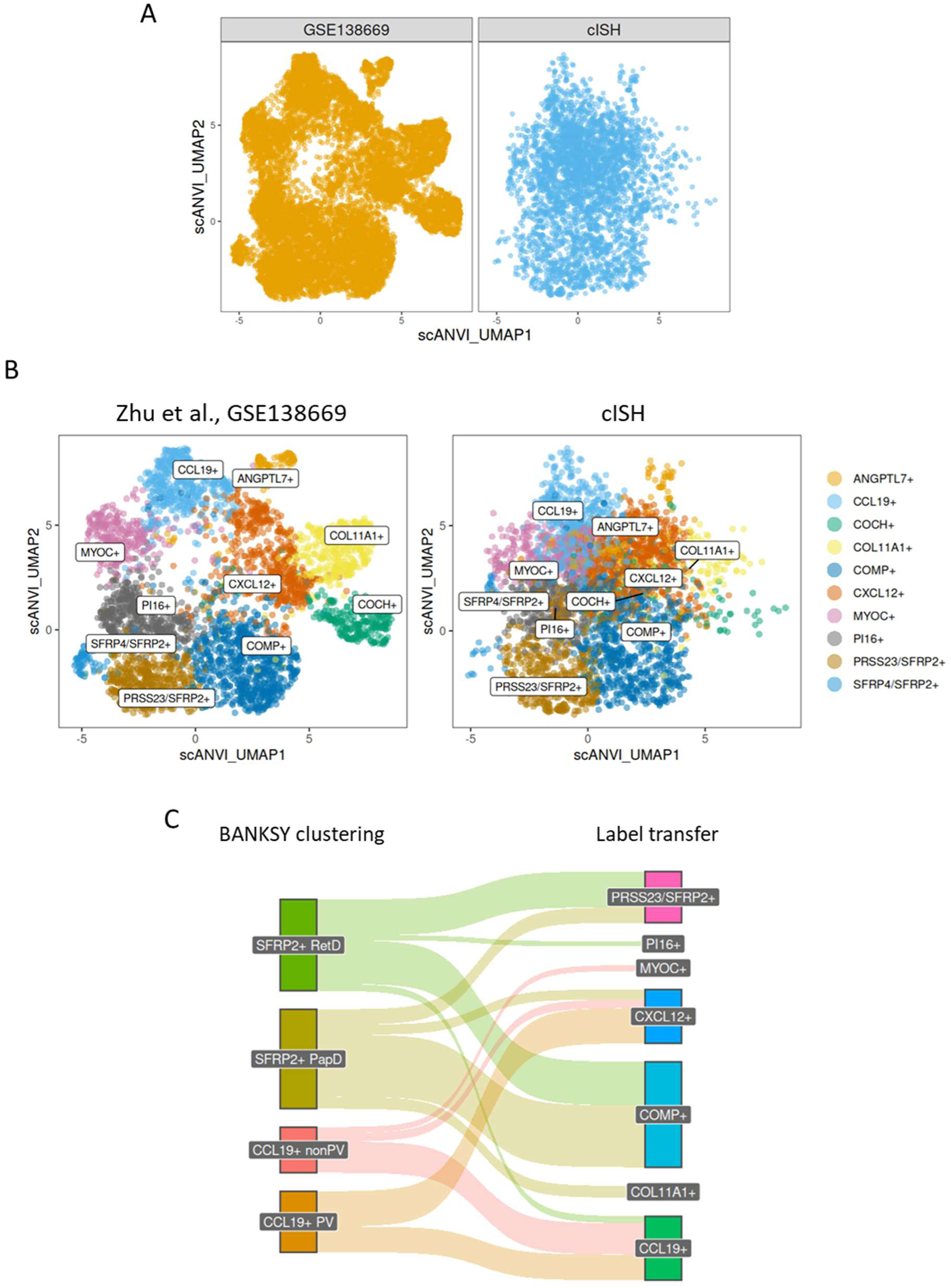
scANVI-based integration of fibroblasts from the cISH dataset with the scRNAseq GSE138669. (A) UMAP plots showing the fibroblasts from our cISH dataset and the scRNA-seq dataset retrieved from Zhu et al. (GEO accession number GSE138669) in an integrated latent space computed by scANVI. (B) UMAP plots showing the fibroblast subsets from the scRNA-seq dataset and the labels transferred to our cISH dataset by scANVI. (C) SANKY plots illustrating the correspondence between the BANKSY-based annotations of spatially segregated fibroblast subsets, on the left side and the annotations by label transfer, on the right side. UMAP, Uniform Manifold Approximation and Projection; GEO, gene expression omnibus; scANVI, single-cell ANnotation using Variational Inference. BANKSY, Building Aggregates with a Neighborhood Kernel and Spatial Yardstick.

**Figure S5:**
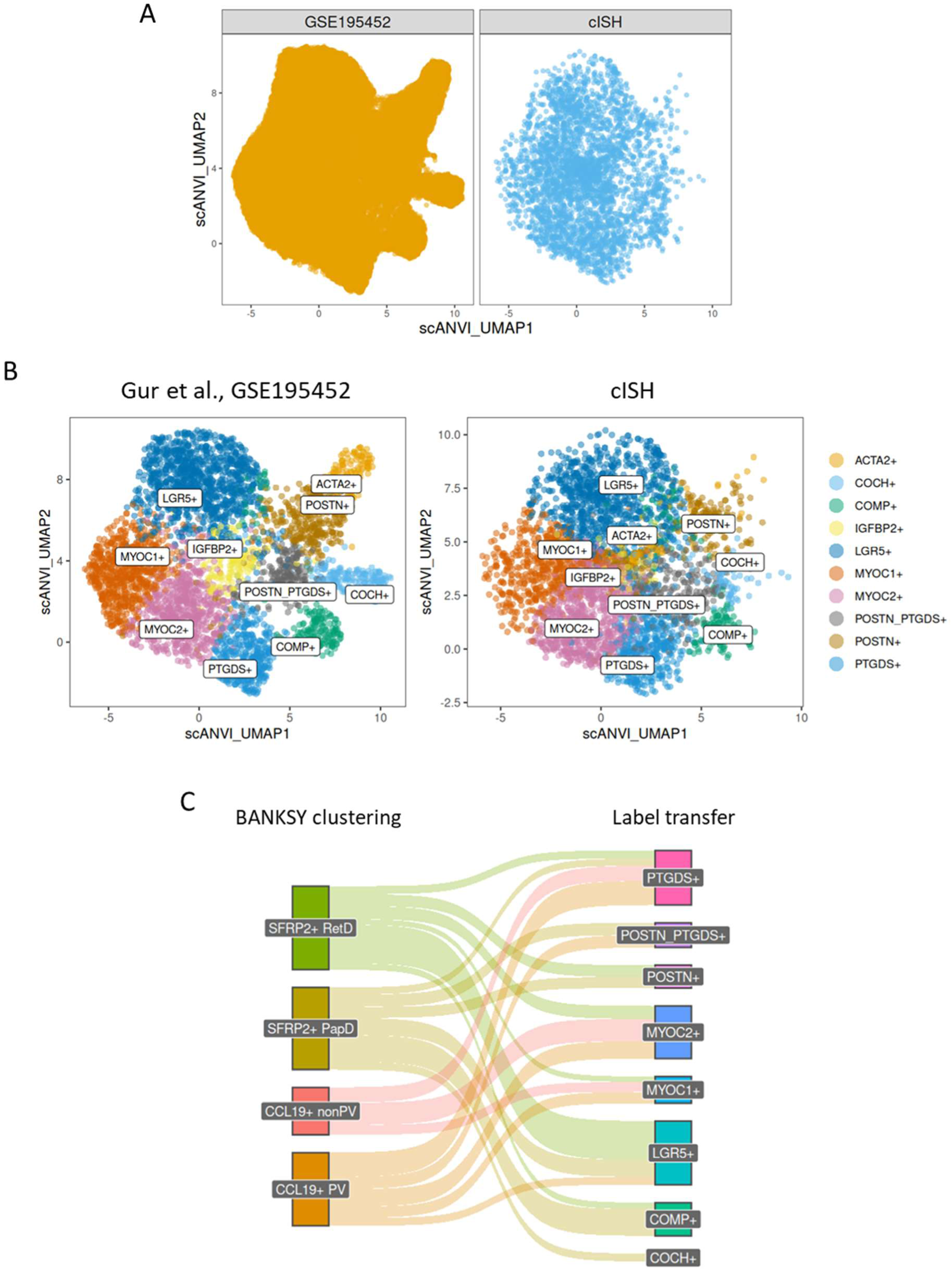
scANVI-based integration of fibroblasts from the cISH dataset with the scRNAseq GSE195452. (A) UMAP plots showing the immune cells from our cISH dataset and the scRNA-seq dataset retrieved from Gur et al. (GEO accession number GSE195452) in an integrated latent space computed by scANVI. (B) UMAP plots showing the fibroblast subsets from the scRNA-seq dataset and the labels transferred to our cISH dataset by scANVI. (C) SANKY plots illustrating the correspondence between the BANKSY-based annotations of spatially segregated fibroblast subsets, on the left side and the annotations by label transfer, on the right side. UMAP, Uniform Manifold Approximation and Projection; GEO, gene expression omnibus; scANVI, single-cell ANnotation using Variational Inference. BANKSY, Building Aggregates with a Neighborhood Kernel and Spatial Yardstick.

**Figure S6:**
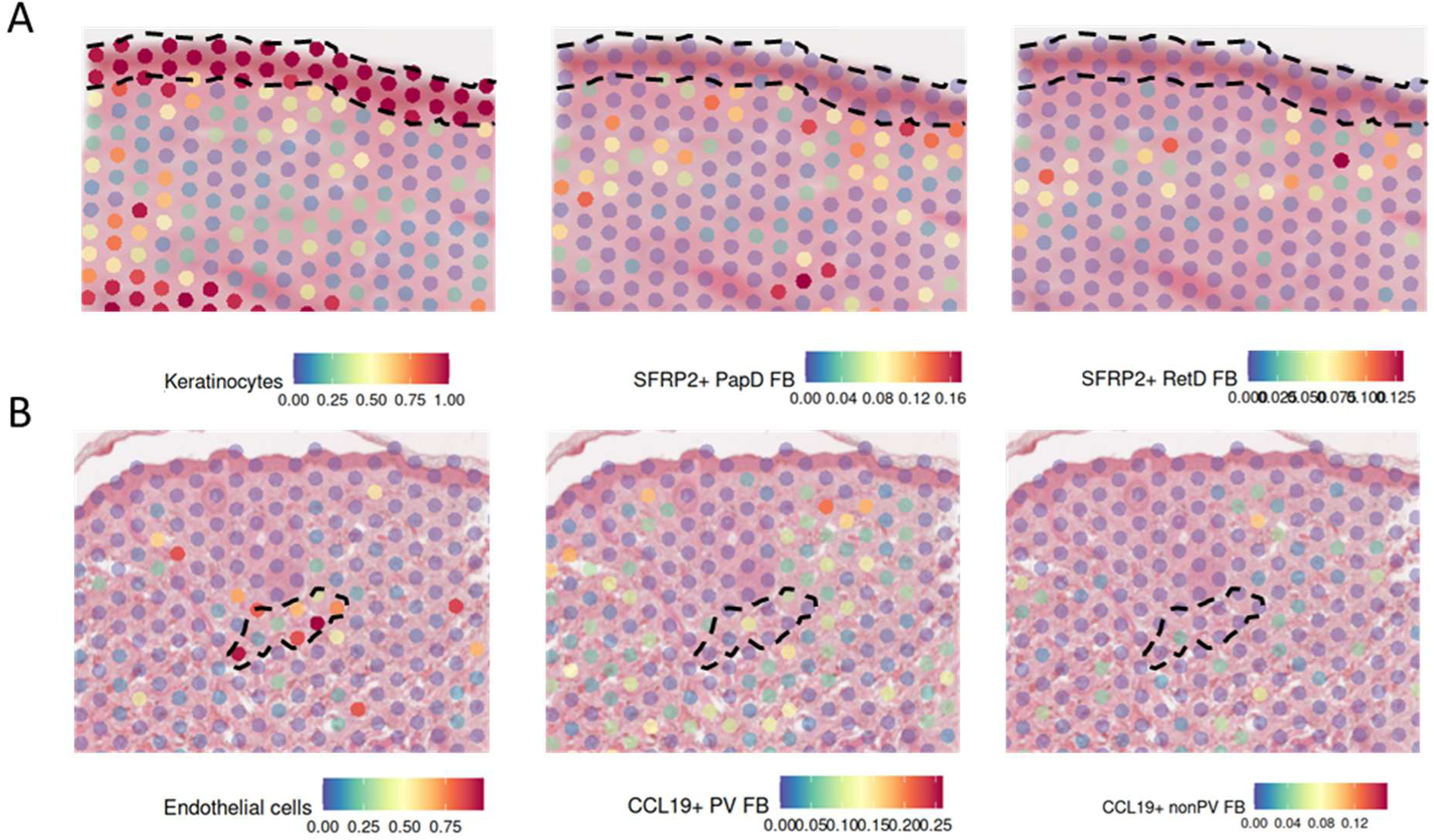
Reference mapping of BANKSY-identified clusters on an independent spatial sequencing dataset. Representative images of predicted cell abundance of keratinocytes with SFRP2+ fibroblasts (A) and endothelial cells with CCL19+ fibroblasts (B) by Seurat anchor-based label transfer from cISH dataset to Visium spatial sequencing dataset of SSc skin. BANKSY, Building Aggregates with a Neighborhood Kernel and Spatial Yardstick.

**Figure S7:**
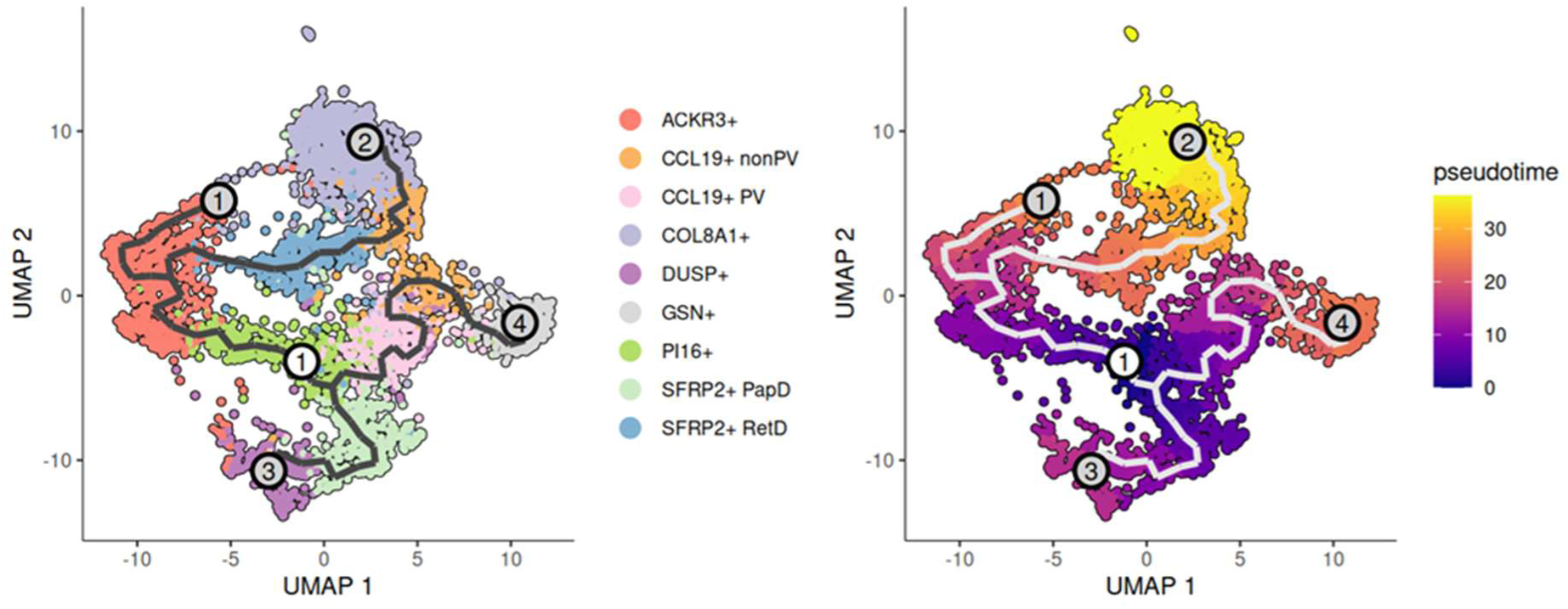
Differentiation trajectories of fibroblast subsets identified by spatially informed phenotyping UMAP plots showing the cell differentiation routes identified by trajectory inference using Monocle, with PI16+ FB as root. UMAP, Uniform Manifold Approximation and Projection.

**Figure S8:**
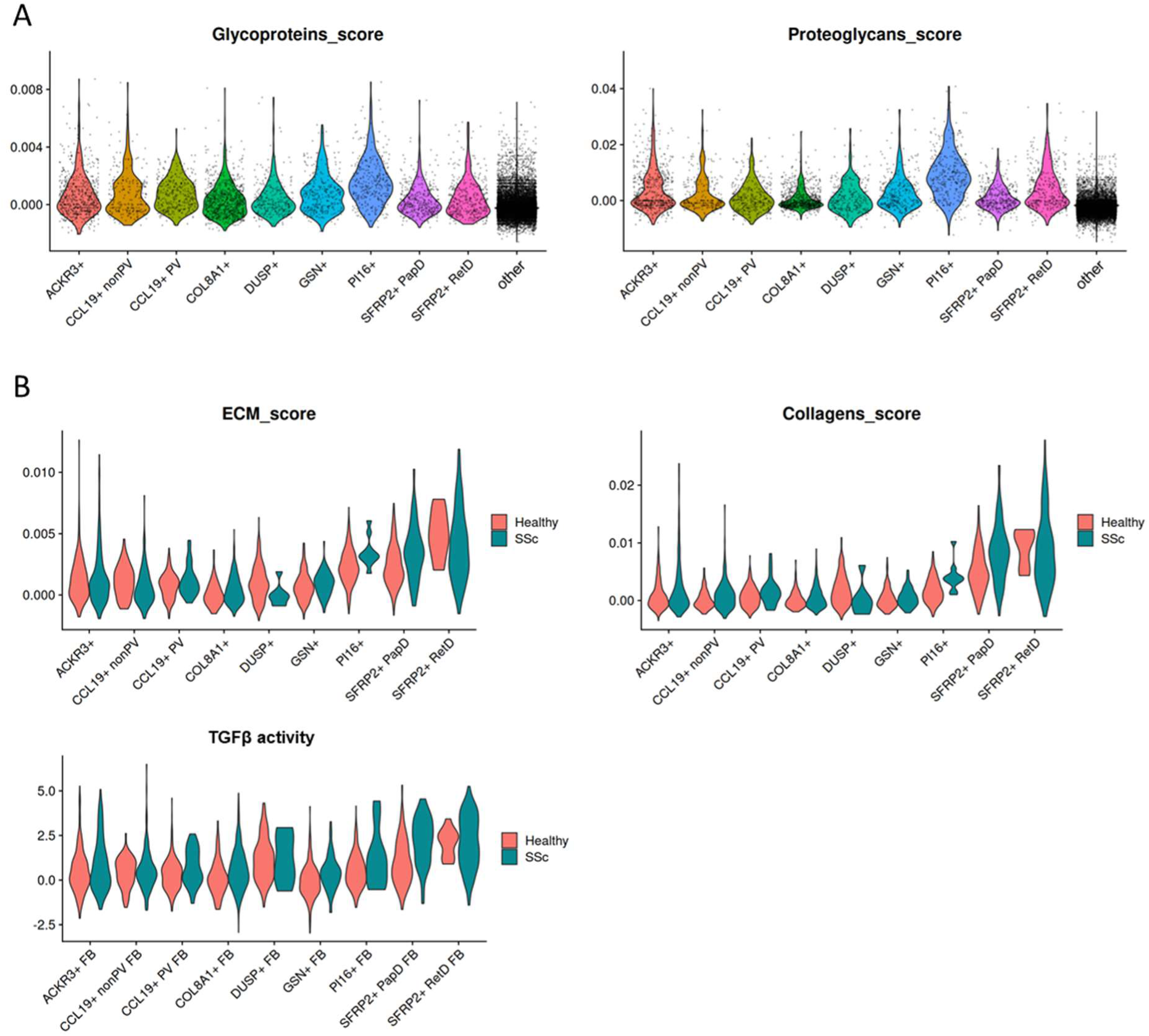
ECM module scores and TGFβ activity across fibroblast populations in healthy and SSc. (A) Violin plots showing the expression scores of glycoprotein and proteoglycan components obtained from the Matrisome project across fibroblast populations. Non-fibroblasts were labeled as ‘other’ in the violin plots. (B) Violin plots showing the expression scores of collagen component, the entire core Matrisome components (marked as ECM score), and TGFβ activities across fibroblast populations in healthy and SSc skin.

**Figure S9:**
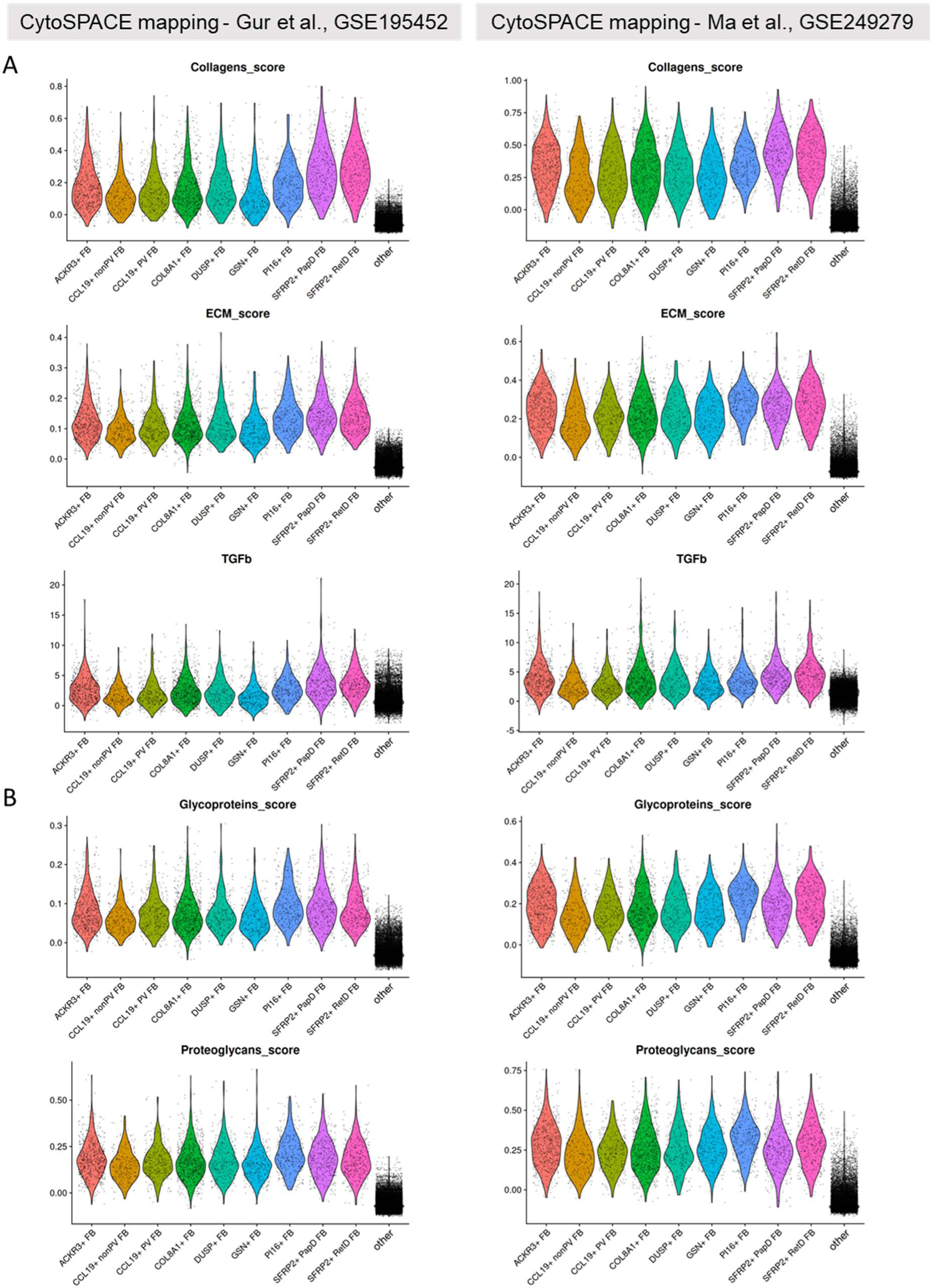
ECM module score and TGFβ activity with CytoSPACE-imputed expression. (A) Violin plots showing the expression scores of collagen component, the entire core Matrisome components (marked as ECM score), and the TGFβ activities across fibroblast populations from imputed expression data computed by CytoSPACE. (B) Violin plots showing the expression scores of glycoproteins and proteoglycans across fibroblast populations from the imputed expression by CytoSPACE. Non-fibroblasts were labeled as ‘other’ in the violin plots.

**Figure S10:**
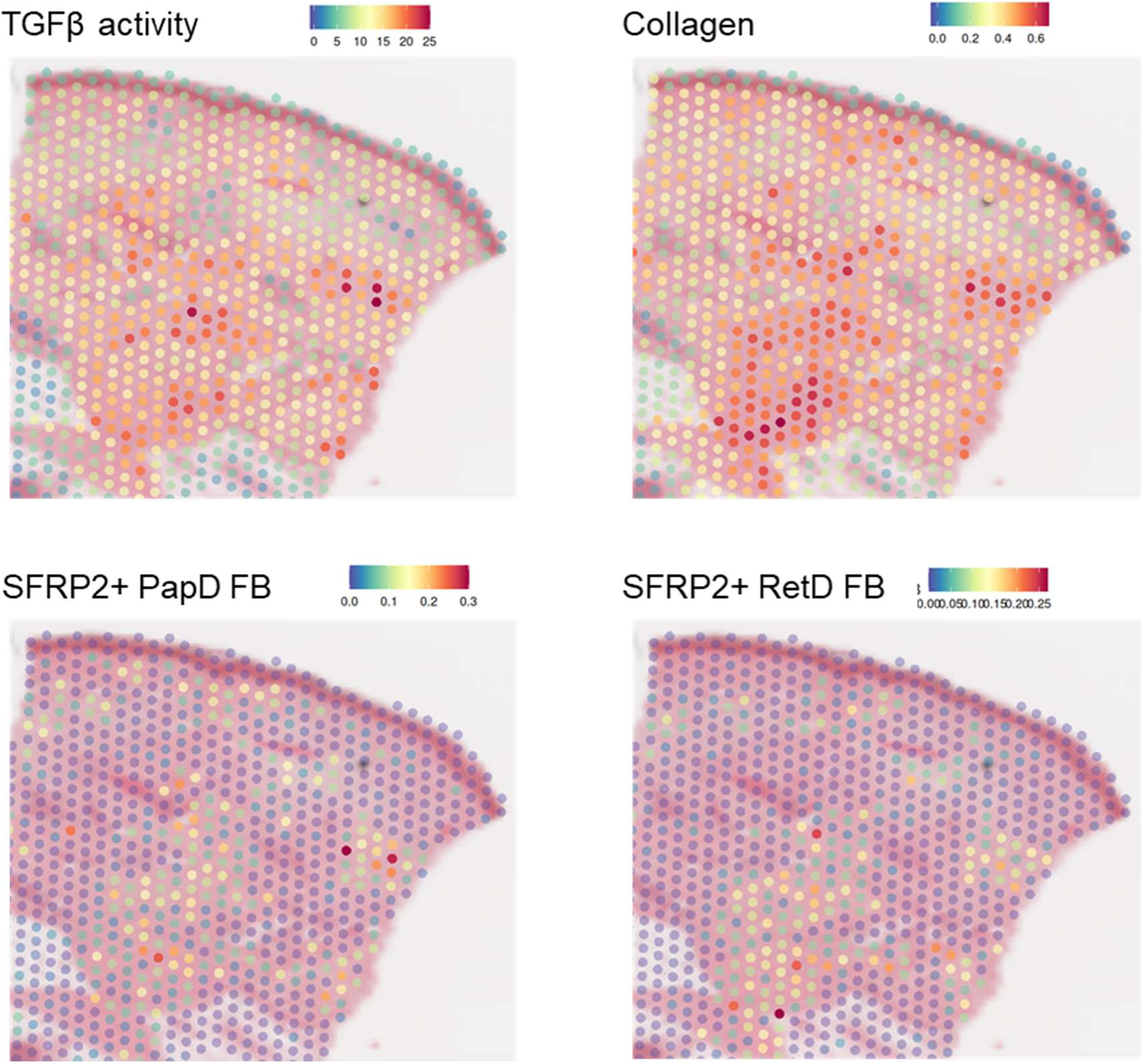
TGFβ activity and collagen score with BANKSY-identified SFRP2+ fibroblasts on an independent spatial sequencing dataset. Representative images showing the TGFβ activities, collagen expression scores, and the abundance of SFRP2+ fibroblasts identified by BANKSY from the cISH dataset. The TGFβ activity and collagen score were computed directly on the expression from the Visium spatial sequencing dataset. The abundance of the BANKSY clusters was obtained from Seurat anchor-based reference mapping. BANKSY, Building Aggregates with a Neighborhood Kernel and Spatial Yardstick.

**Figure S11:**
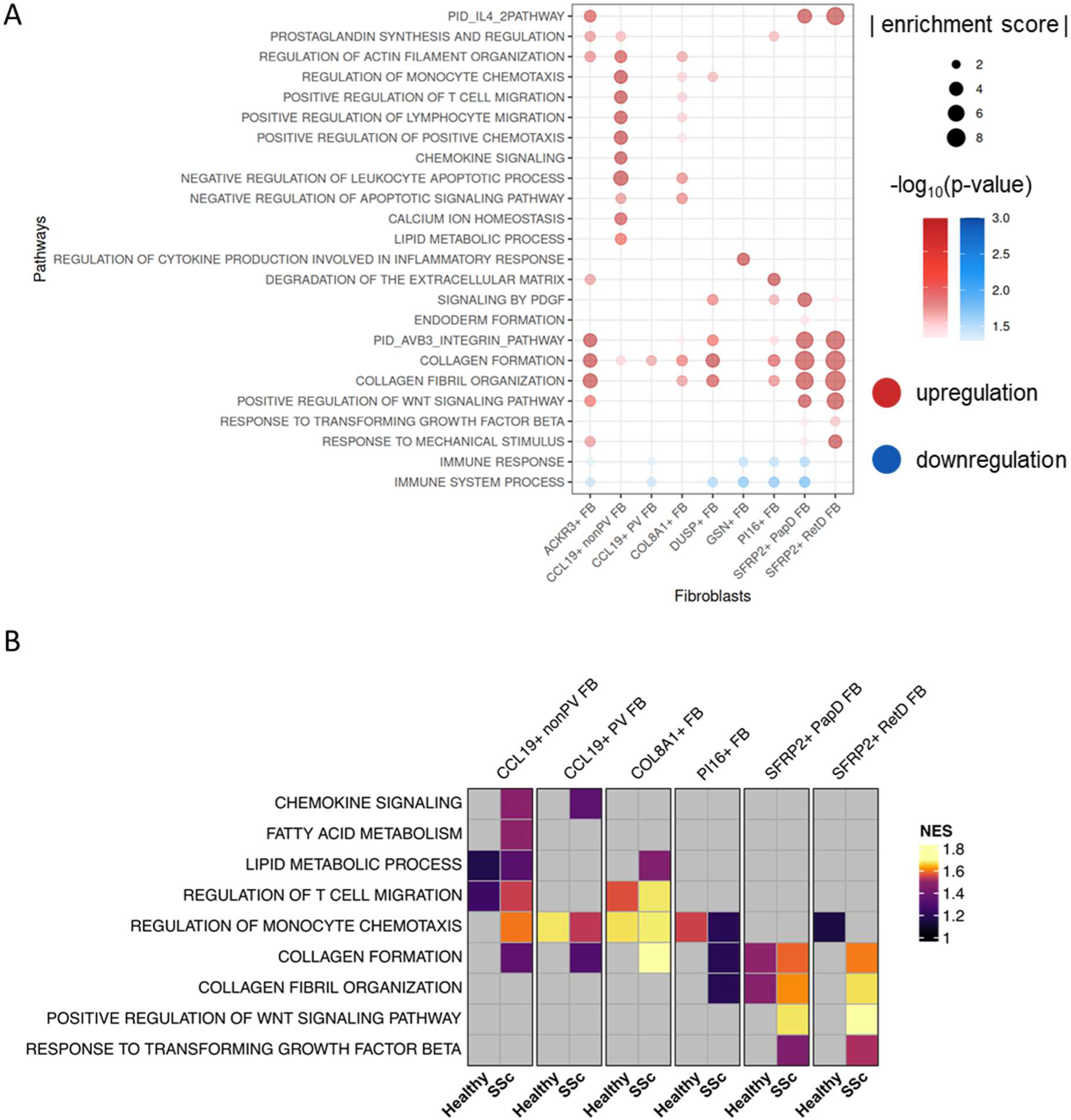
Distinct biological functions of fibroblast populations. Dot plots showing the enriched functional pathway across fibroblast populations by ULM(A) and FGSEA (B). The pathway enrichment analysis was performed using decoupleR R package using the pseudo-bulk expression matrix that contained averaged expression for each fibroblast subpopulation and group. Pathways that did not reach statistical significance (p-value ≥ 0.05) were excluded from the plots. The size of the dot represents NES (A) or absolute enrichment score (B). Up- and downregulation were defined by significant positive and negative NES or enrichment score, respectively. FGSEA, fast gene set enrichment analysis; ULM, univariate linear model; NES, normalized enrichment score.

**Figure S12:**
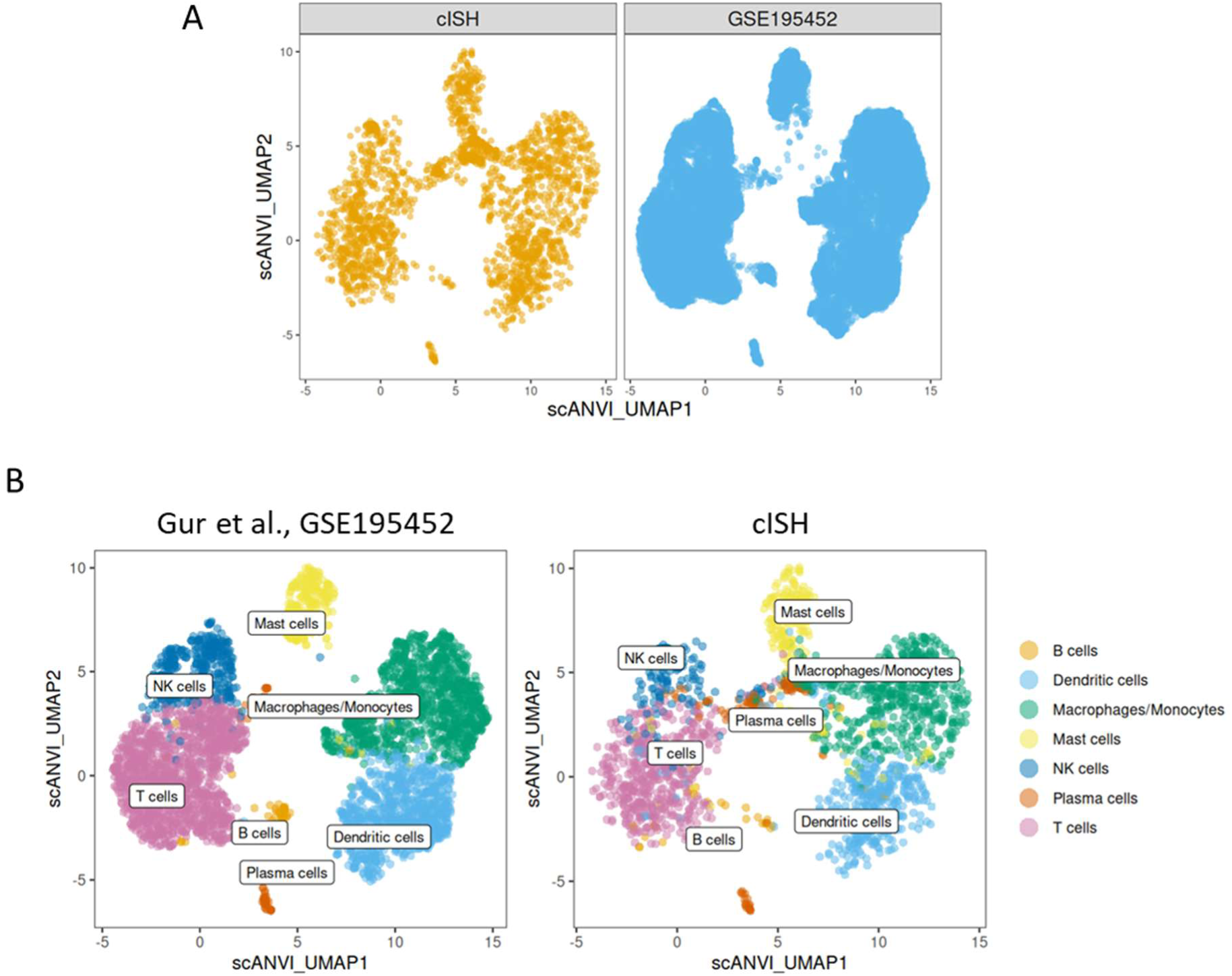
Annotation of immune cells through scANVI-based integration and label transfer. (A) UMAP plots showing the immune cells from our cISH dataset and the scRNA-seq dataset retrieved from Gur et al. (GEO accession number GSE195452) in an integrated latent space computed by scANVI. (B) UMAP plots showing the immune cell populations from the scRNA-seq dataset and the labels transferred to our cISH dataset by scANVI. UMAP, Uniform Manifold Approximation and Projection; GEO, gene expression omnibus; scANVI, single-cell ANnotation using Variational Inference.

**Figure S13:**
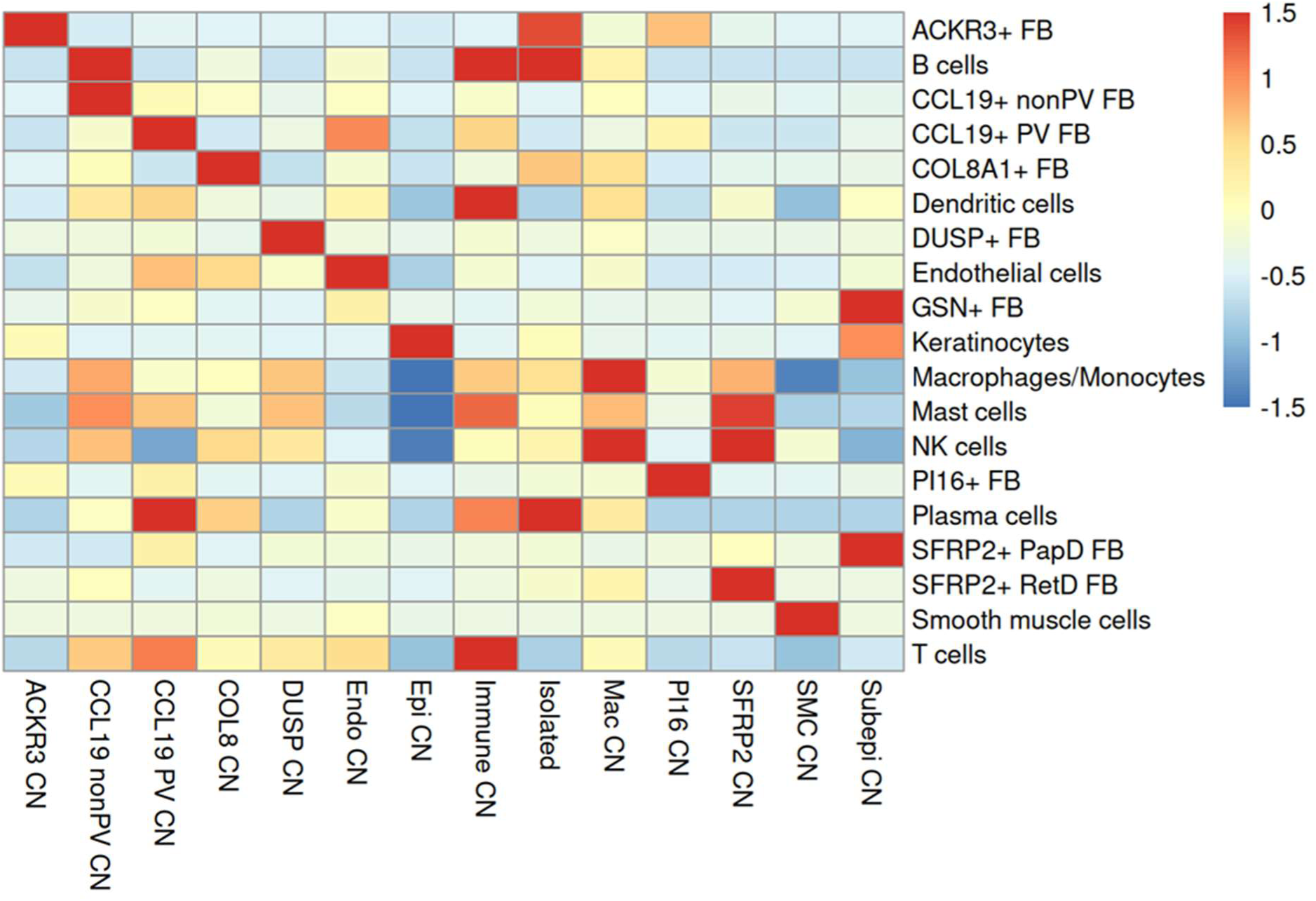
Cellular neighborhoods detected based on the composition of neighboring cell types. Heatmap showing the composition of 14 CNs identified by k-means clustering on the neighboring cell types for each cell. The composition was normalized for each cell type for visualization purposes. CN, cellular neighborhood.

**Figure S14:**
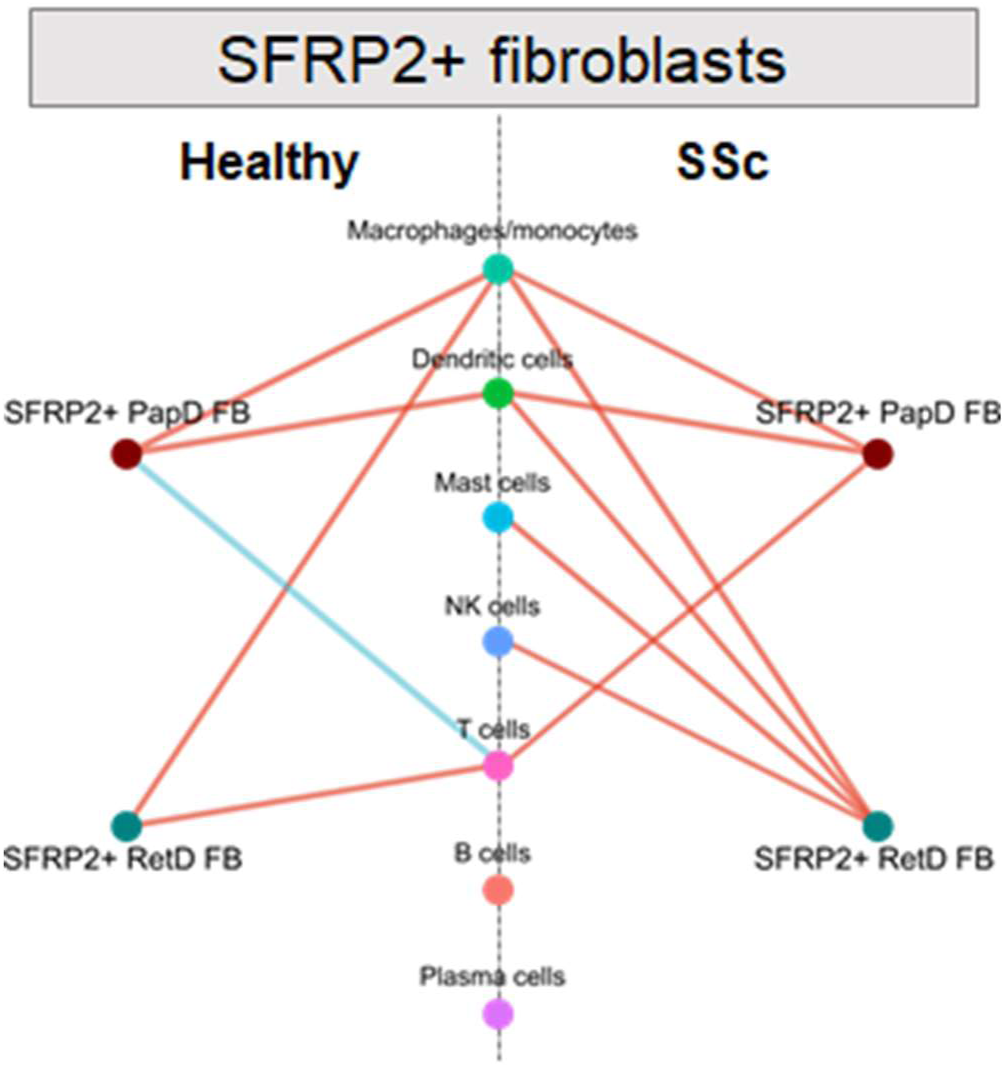
Pairwise interaction analysis between immune cells and SFRP2+ fibroblasts. Network visualization of immune interaction with two SFRP2+ fibroblast populations detected by spatial clustering in healthy and SSc skin based on spatial neighboring pairs by pairwise interaction analysis. The cellular interactions are shown as the lines in the plot. The color indicates cellular avoidance (blue) and attraction (red). The width of the lines indicates the proportion of samples with statistically significant interactions found in the indicated group. The statistical significance (p < 0.05) was determined by permutation test which shuffles the cell labels to build a random spatial distribution of the cells.

**Figure S15:**
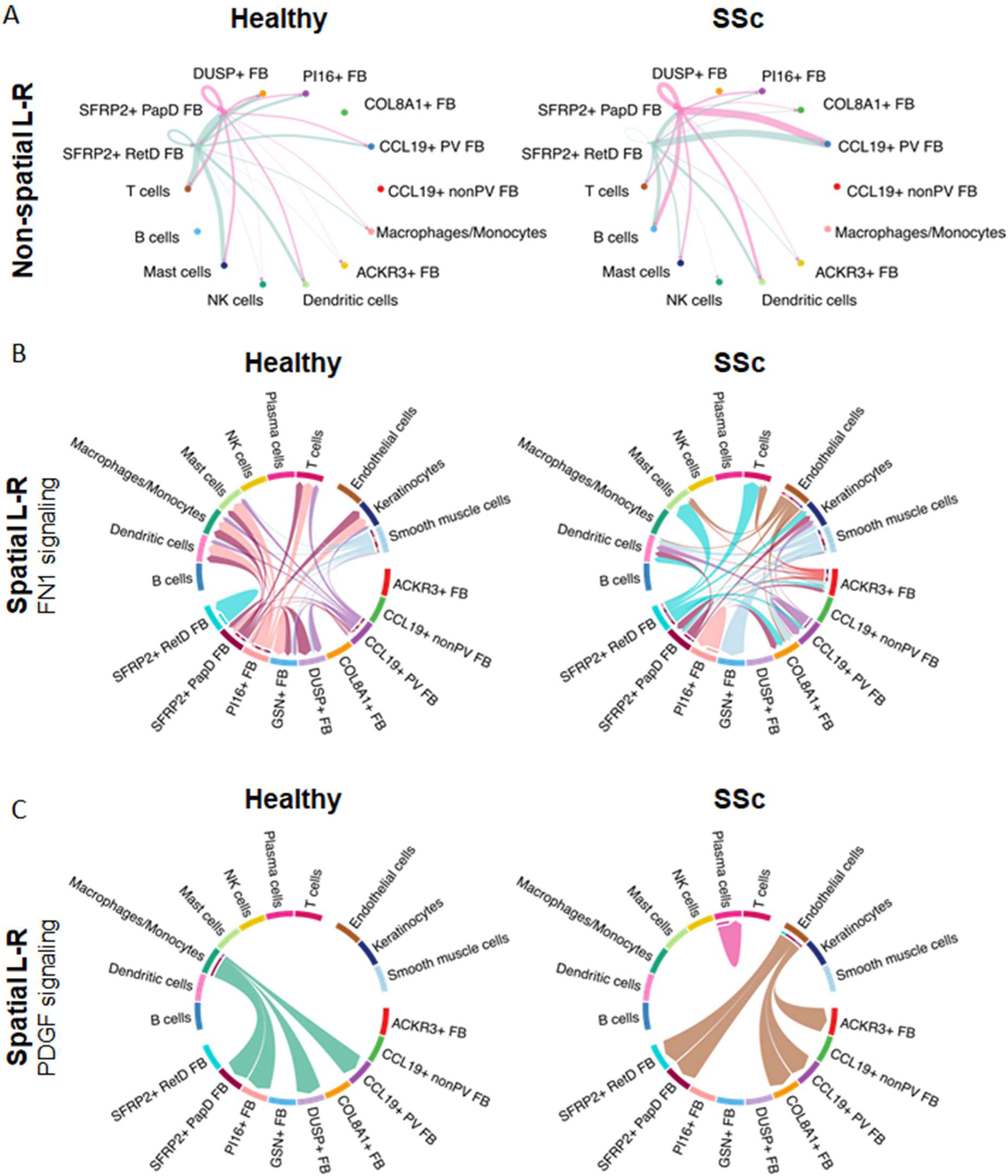
Changes in cellular communication in SSc detected by ligand-receptor analysis. (A) Circle plots showing the interaction strength for the signals coming from SFRP2+ fibroblasts in healthy and SSc skin by non-spatial L-R analysis using CellChat R package. (B-C) Chord diagram showing FN1 signaling (B) or PDGF signaling (C) detected by spatial L-R analyses in healthy and SSc skin. L-R analysis, ligand-receptor analysis.

**Figure S16:**
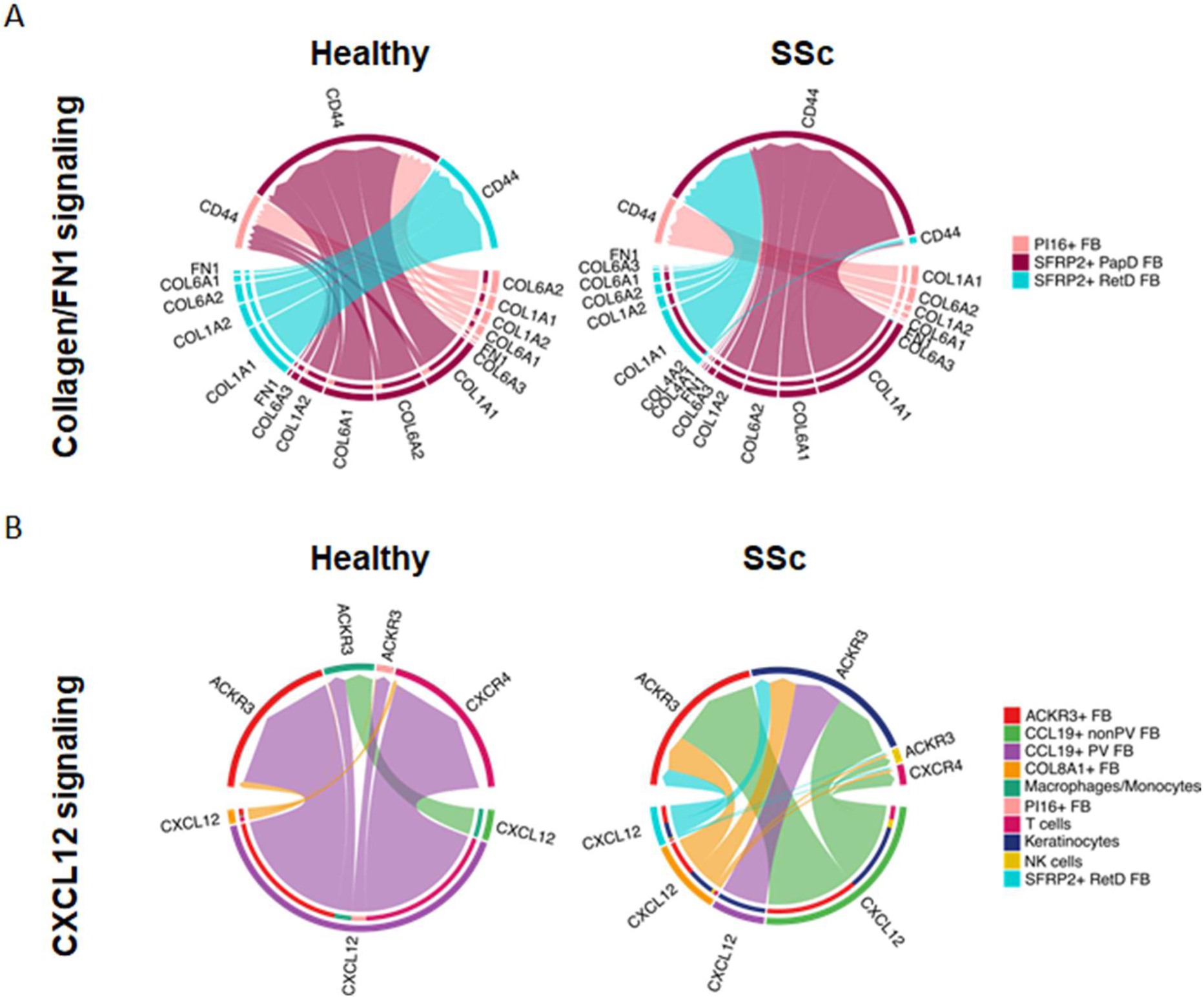
Key molecules of ECM- and chemokine signaling in SSc identified by spatial L-R analysis. (A) The self ECM-sensing ability shown by the chord diagram of L-R analysis on collagen and FN1 signaling among SFRP2+ fibroblasts and PI16+ FB. (B) Chord diagrams of L-R interaction of CXCL12 signaling from CCL19+ fibroblasts, COL8A1+ FB, and SFRP2+ RetD FB in healthy and SSc skin. The L-R pairs are indicated on the chord diagrams. The inner bars in the diagram indicate the interaction strength towards the indicated target cells. ECM, extracellular matrix; L-R, ligand-receptor.

**Figure S17:**
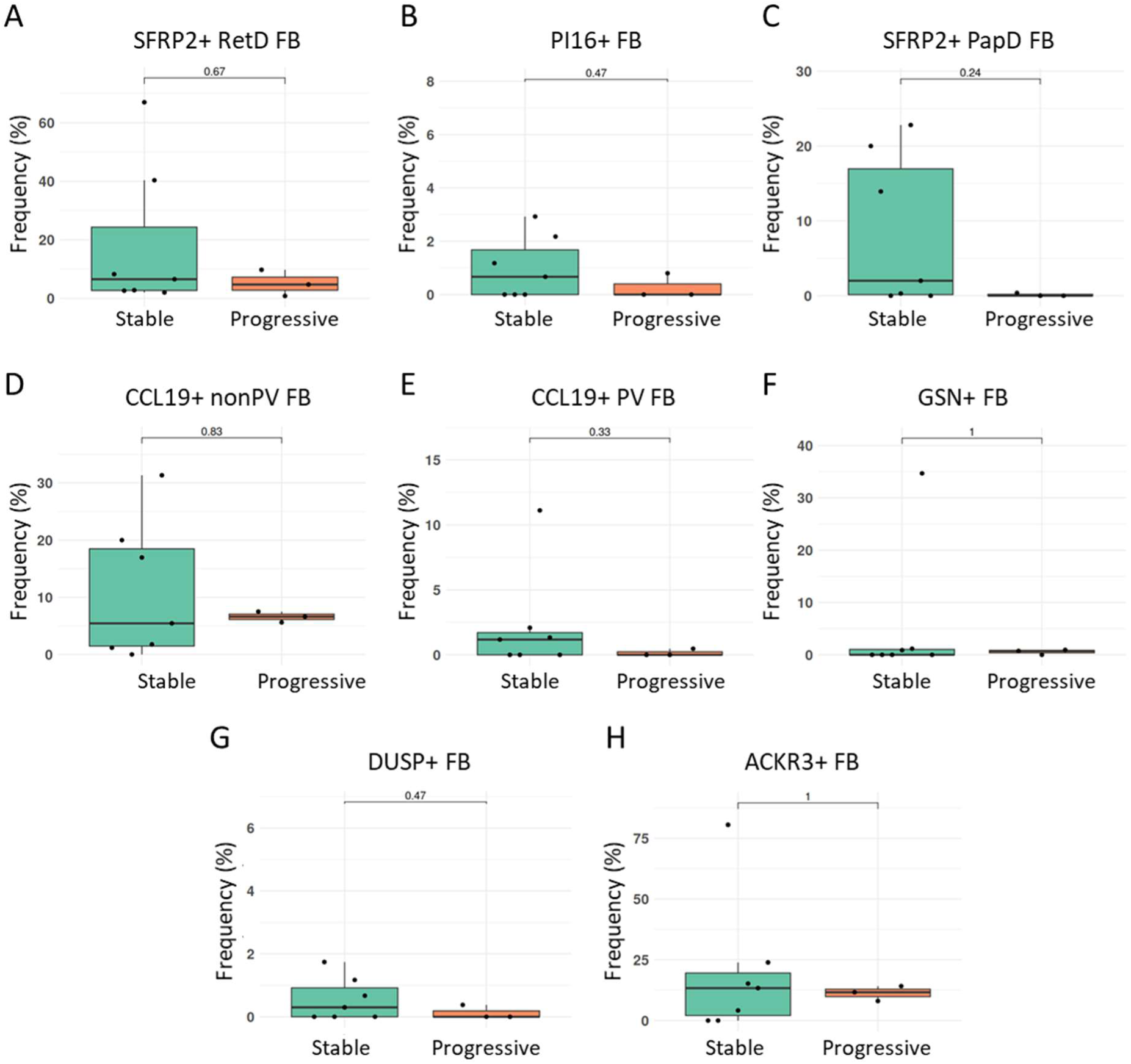
Frequencies of fibroblast subsets other than COL8A1+ FB in progressive skin fibrosis. (A-H) Frequencies of SFRP2+ RetD FB (A), of PI16+ FB (B), of SFRP2+ PapD FB (C), of CCL19+ nonPV FB (D), of CCL19+ PV FB (E), of GSN+ FB (F), of DUSP+ FB (G) and of ACKR3+ FB (H) among all dermal cells in patients with progressive or stable skin fibrosis. (I-L) Frequencies of COL8A1+ FB in spatial proximity to T cells (I), to dendritic cells (J), to NK cells (K) or to plasma cells (L) among all dermal cells. FB, fibroblasts; PapD, papillary dermis; RetD, reticular dermis; PV, perivascular.

**Figure S18:**
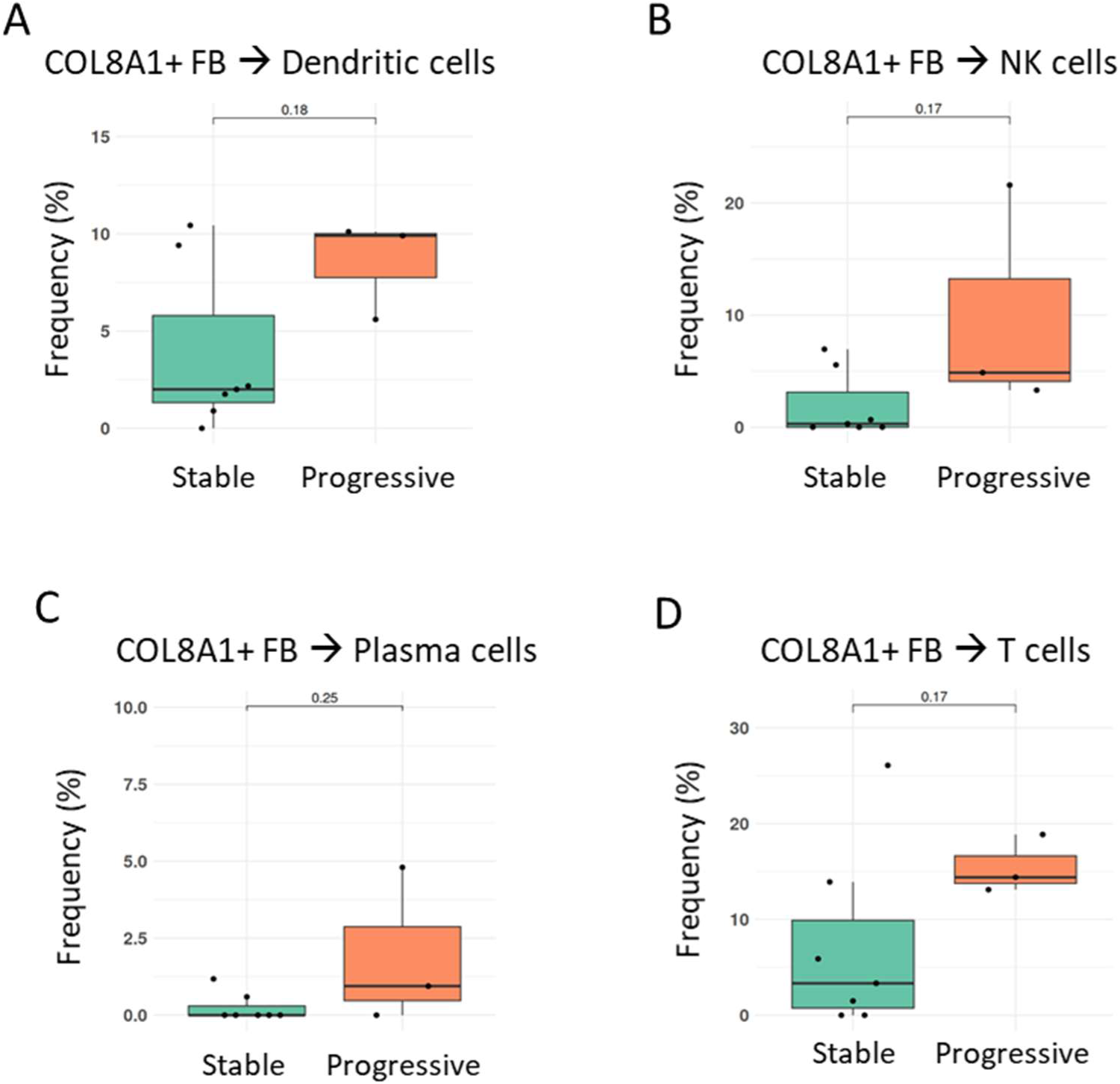
Frequencies of COL8A1+ FB in spatial proximity to specific immune cell subsets (other than those shown in main figure 7) in progressive skin fibrosis. (A-D) Frequencies of COL8A1+ FB in spatial proximity to dendritic cells (A), to NK cells (B), to plasma cells (C) or to T cells (D), among all dermal cells. FB, fibroblasts; NK, natural killer.

**Figure S19:**
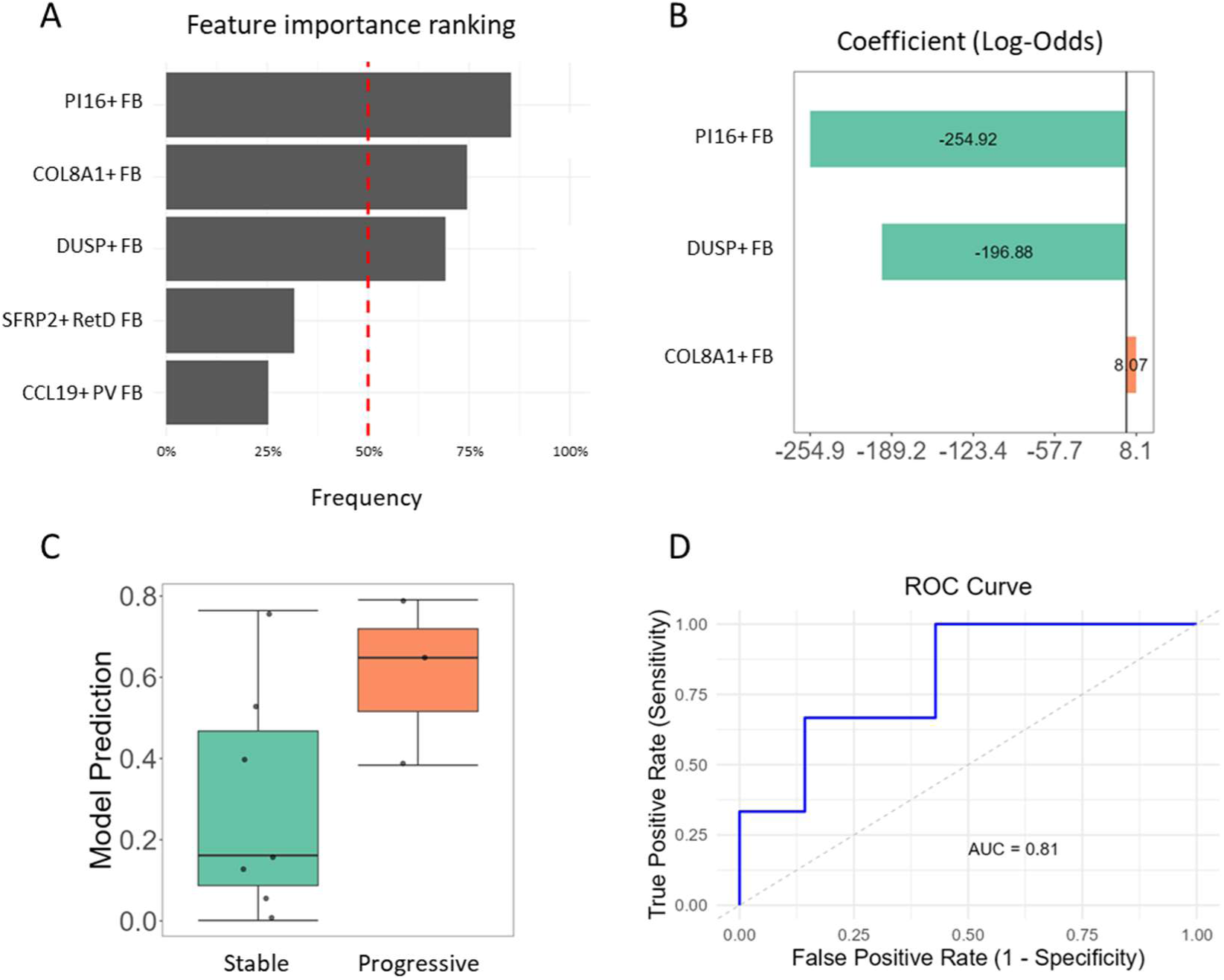
Prediction of progression of skin fibrosis in SSc by fibroblast frequencies (A) Top highly ranked features predictive of progression of skin fibrosis after bootstrapping with LASSO regression starting from the frequencies of all COL8A1+ FB, of SFRP2+ RetD FB, of PI16+ FB, of DUSP FB+ and of CCL19+ PV FB as predictors. (B) Increased frequencies of COL8A1+ fibroblasts (FB) were associated with a higher likelihood of progressive skin fibrosis, whereas elevated frequencies of PI16+ FB and DUSP+ FB were linked to a decreased likelihood of fibrosis progression. (C) Model prediction results for each donor. (D) ROC curve and its AUC for evaluation of model performance. FB, fibroblasts; RetD, reticular dermis; LASSO, least absolute shrinkage and selection operator; ROC, receiver operating characteristic; AUC, area under the curve.

## Supplementary tables

**Table S1:**
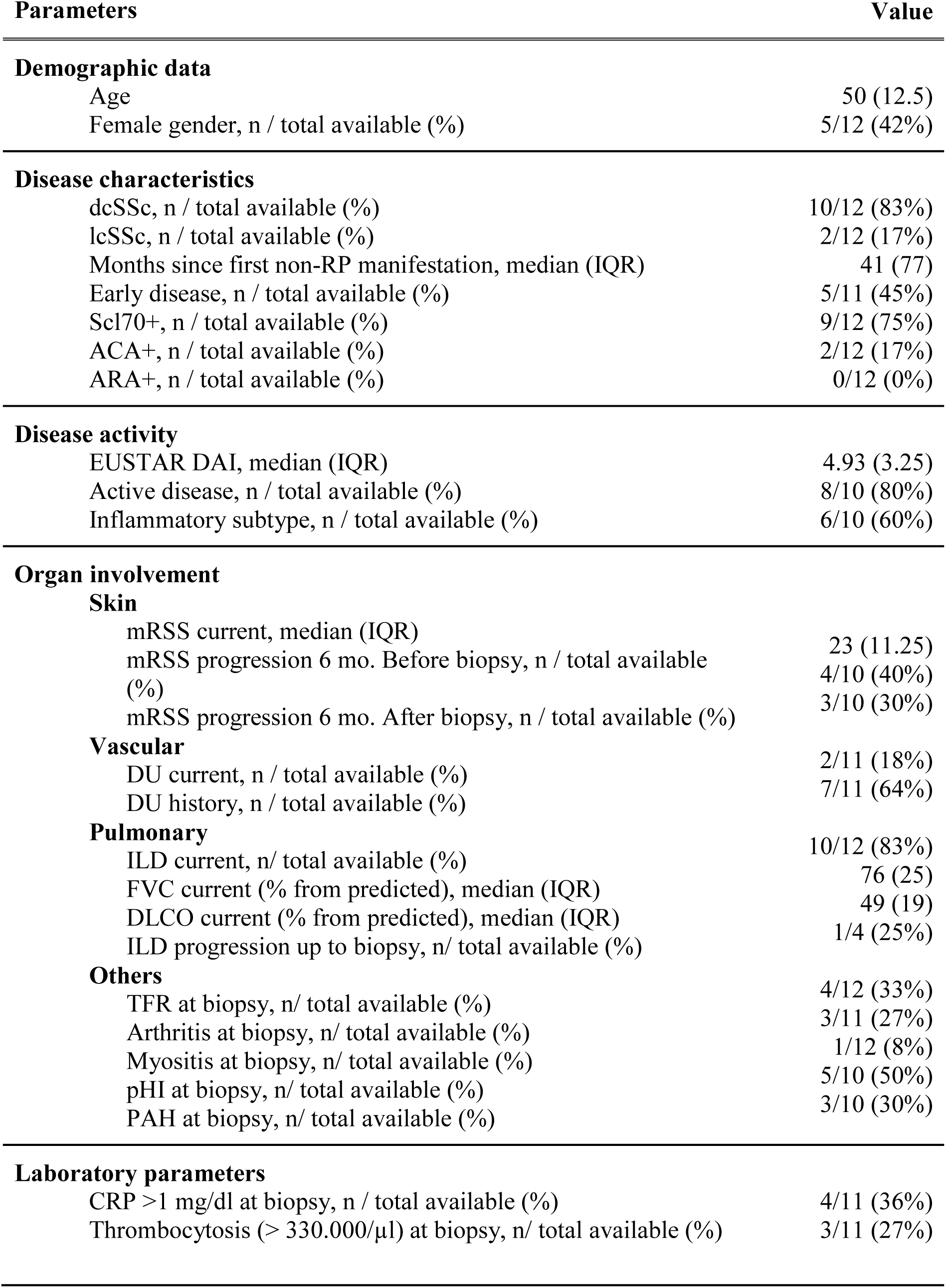
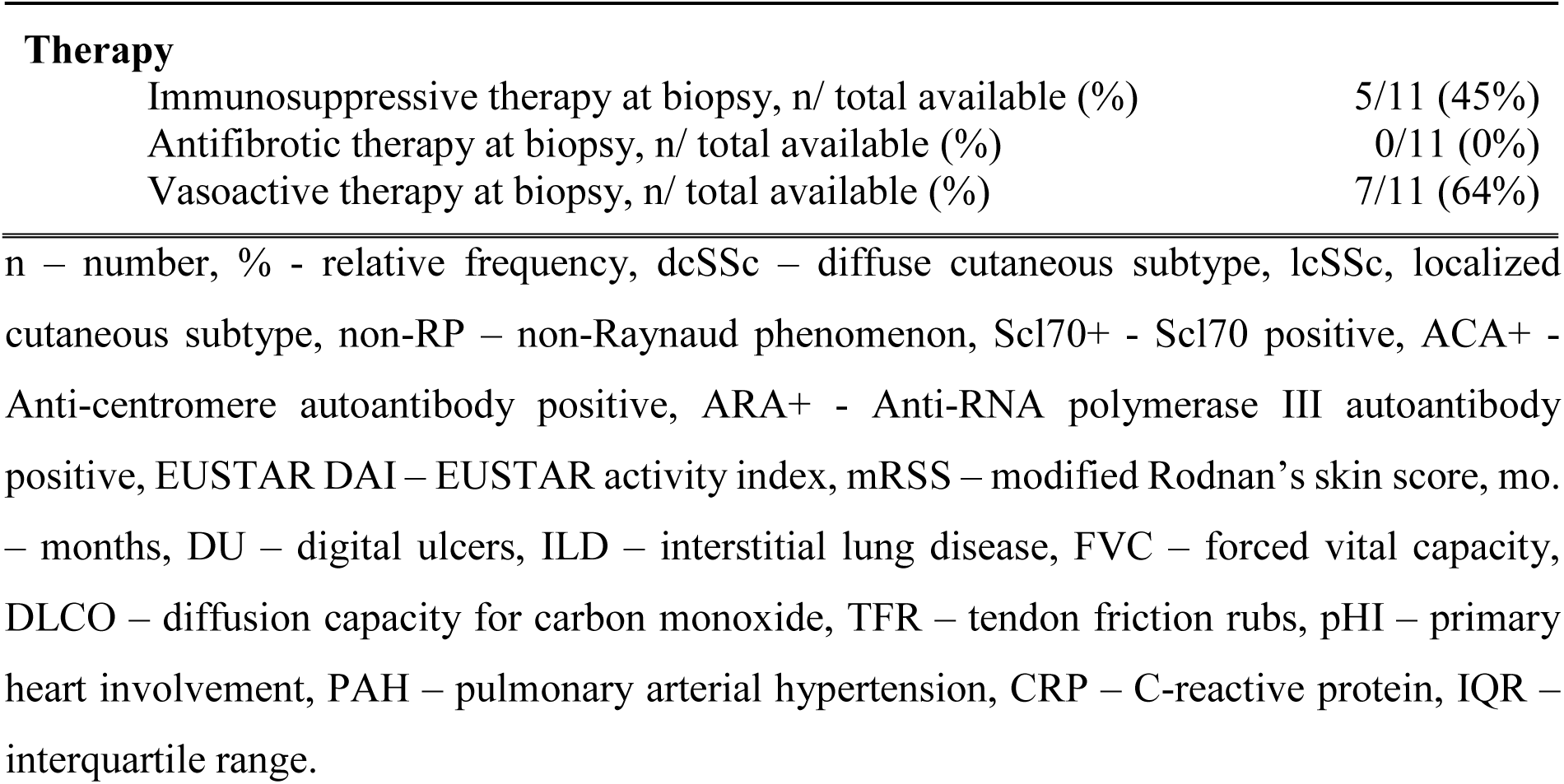
Clinical features of the SSc donors.

| Parameters | Value |
| --- | --- |
| **Demographic data** |  |
| Age | 50 (12.5) |
| Female gender, n / total available (%) | 5/12 (42%) |
| **Disease characteristics** |  |
| dcSSc, n / total available (%) | 10/12 (83%) |
| lcSSc, n / total available (%) | 2/12 (17%) |
| Months since first non-RP manifestation, median (IQR) | 41 (77) |
| Early disease, n / total available (%) | 5/11 (45%) |
| Scl70+, n / total available (%) | 9/12 (75%) |
| ACA+, n / total available (%) | 2/12 (17%) |
| ARA+, n / total available (%) | 0/12 (0%) |
| **Disease activity** |  |
| EUSTAR DAI, median (IQR) | 4.93 (3.25) |
| Active disease, n / total available (%) | 8/10 (80%) |
| Inflammatory subtype, n / total available (%) | 6/10 (60%) |
| **Organ involvement** |  |
| **Skin** |  |
| mRSS current, median (IQR) | 23 (11.25) |
| mRSS progression 6 mo. Before biopsy, n / total available (%) | 4/10 (40%) |
| mRSS progression 6 mo. After biopsy, n / total available (%) | 3/10 (30%) |
| **Vascular** |  |
| DU current, n / total available (%) | 2/11 (18%) |
| DU history, n / total available (%) | 7/11 (64%) |
| **Pulmonary** |  |
| ILD current, n/ total available (%) | 10/12 (83%) |
| FVC current (% from predicted), median (IQR) | 76 (25) |
| DLCO current (% from predicted), median (IQR) | 49 (19) |
| ILD progression up to biopsy, n/ total available (%) | 1/4 (25%) |
| **Others** |  |
| TFR at biopsy, n/ total available (%) | 4/12 (33%) |
| Arthritis at biopsy, n/ total available (%) | 3/11 (27%) |
| Myositis at biopsy, n/ total available (%) | 1/12 (8%) |
| pHI at biopsy, n/ total available (%) | 5/10 (50%) |
| PAH at biopsy, n/ total available (%) | 3/10 (30%) |
| **Laboratory parameters** |  |
| CRP >1 mg/dl at biopsy, n / total available (%) | 4/11 (36%) |
| Thrombocytosis (> 330.000/μl) at biopsy, n/ total available (%) | 3/11 (27%) |
| **Therapy** |  |
| Immunosuppressive therapy at biopsy, n/ total available (%) | 5/11 (45%) |
| Antifibrotic therapy at biopsy, n/ total available (%) | 0/11 (0%) |
| Vasoactive therapy at biopsy, n/ total available (%) | 7/11 (64%) |

n – number, % - relative frequency, dcSSc – diffuse cutaneous subtype, lcSSc, localized cutaneous subtype, non-RP – non-Raynaud phenomenon, Scl70+ - Scl70 positive, ACA+ - Anti-centromere autoantibody positive, ARA+ - Anti-RNA polymerase III autoantibody positive, EUSTAR DAI – EUSTAR activity index, mRSS – modified Rodnan’s skin score, mo. – months, DU – digital ulcers, ILD – interstitial lung disease, FVC – forced vital capacity, DLCO – diffusion capacity for carbon monoxide, TFR – tendon friction rubs, pHI – primary heart involvement, PAH – pulmonary arterial hypertension, CRP – C-reactive protein, IQR – interquartile range.

